# Slippery sequences stall the 26S proteasome at multiple points along the translocation pathway

**DOI:** 10.1101/2024.03.14.584990

**Authors:** Edwin R. Ragwan, Faith M. Kisker, Amelia R. Morning, Kaya R. Weiser, Athena V. Lago, Daniel A. Kraut

## Abstract

In eukaryotes, the ubiquitin-proteasome system is responsible for intracellular protein degradation. Proteins tagged with ubiquitin are recognized by ubiquitin receptors on the 19S regulatory particle (RP) of the 26S proteasome, unfolded, routed through the translocation channel of the RP, and are then degraded in the 20S core particle (CP). Aromatic paddles on the pore-1 loops of the RP’s Rpt subunits grip the substrate and pull folded domains into the channel, thereby unfolding them. The sequence that the aromatic paddles grip while unfolding a substrate is therefore expected to influence the extent of unfolding, and low complexity sequences have been shown to interfere with grip. However, the detailed spatial requirements for grip while unfolding proteins, particularly from the N-terminus, remain unknown. We determined how the location of glycine-rich tracts relative to a folded domain impairs unfolding. We find that, in contrast to a previous report, inserting glycine-rich sequences closer to the folded domain reduced unfolding ability more than positioning them further away. Locations that have the biggest effect on unfolding map onto the regions where the aromatic paddles are predicted to interact with the substrate. Effects on unfolding from locations up to 67 amino acids away from the folded domain suggest that there are additional interactions between the substrate and the proteasome beyond the aromatic paddles that facilitate translocation of the substrate. In sum, this study deepens understanding of the mechanical interactions within the substrate channel by mapping the spacing of interactions between the substrate and the proteasome during unfolding.

**Importance:** The proteasome processively unfolds and degrades target proteins in eukaryotes. However, some substrates are prematurely released, and the resulting partially degraded proteins can cause problems for cells and can be linked to neurodegenerative diseases. In this paper, we use a series of substrates that can stall the proteasome during degradation to probe the translocation pathway substrates must traverse during unfolding. We find that multiple points along the translocation pathway are impacted by these slippery substrates.

## Introduction

The regulation of intracellular proteins is vital for the cell, and a major mode of regulation is the removal of unwanted proteins. Misfolded proteins can become toxic, and many regulatory proteins need to be degraded and recycled after they have served their function.^1–3^ The ubiquitin-proteasome system (UPS) carries out the majority of protein degradation in eukaryotic cells, with targets identified and tagged via ubiquitin ligases and then delivered to the 26S proteasome for degradation.^4,5^ The proteasome is a large barrel-shaped multi-subunit protease made up of a 20S core particle (CP), where degradation occurs, capped by one or two 19S regulatory particles (RP). Typically, a protein destined for degradation is tagged with a polyubiquitin chain that allows it to dock at proteasomal ubiquitin receptors in the RP. An unstructured region on the substrate is then engaged by ATPase motor proteins within the RP.^6,7^ Access to proteolysis is routed through a substrate channel that is too small to allow passage of folded proteins.^8^ Therefore, the substrate needs to be unfolded and translocated into the buried CP where it is degraded into peptide fragments.

The proteasome adopts various conformations ranging from substrate-binding to substrate-degrading states. The structure of the CP remains relatively consistent, however the RP, which forms base and lid subcomplexes, undergoes major conformational changes.^9^ For example, substrate binding leads to the lid subcomplex rotating such that Rpn11, the essential deubiquitinase responsible for removing ubiquitin from the substrate, becomes coaxially aligned above the central processing pore of the base, facilitating translocation-coupled deubiquitination. At the base of the RP, AAA+-ATPase motor proteins form a central pore, made up of a hexamer of Rpt1-6 subunits that becomes aligned with the axial pore of the CP. To become degraded, the outer rings of the degradation chamber, and their N-termini, open up to allow the substrate to be translocated to the proteolytic sites.

The pore-1 loop from each subunit, referred to as the “aromatic paddle”, contributes a tyrosine residue that protrudes into the pore. These tyrosines are arranged in a spiral staircase configuration, which provides a pathway for substrate translocation through the pore. First, an unstructured region of the substrate serves as an initiation site, which is engaged by the pore loops within the axial channel of the RP.^10,11^ ATP hydrolysis provides the force needed to translocate the substrate into the CP. The substrate then moves through the pore in a proposed “hand-over-hand” mechanism where each aromatic paddle moves the substrate toward the CP until the paddles reach the end of that region, disengage, and return to and re-engage with an upstream region of the substrate.^10–12^ An additional pore-2 loop from each ATPase subunit that forms a second staircase near the pore-1 loop has also been implicated in translocation.^12,13^

Although the proteasome is highly processive and has the structural flexibility to accommodate a wide variety of substrates, there are times when substrates are only partially degraded, either intentionally or unintentionally.^14–16^ One example in mammalian cells is the activation of the transcription factor NF-κB, in which the C-terminal inhibitory region of p105 is degraded by the proteasome to form the functional p50 subunit.^15^ A glycine-rich region (GRR) in p105, combined with a nearby stably folded domain, likely serves as a “stop signal” that prevents the proteasome from fully degrading p105.^17^ Other examples of partial proteasomal degradation have been observed in multiple eukaryotic species using other low-complexity regions (Gly-Ala, serine-rich, etc.), and similar phenomena have been observed in proteins partially degraded by bacterial ATP-dependent proteases.^18,19^ Partial proteasomal degradation may also play a role in the accumulation of protein aggregates in neurodegenerative diseases, such as the glutamine-repeat-containing proteins that are characteristic of Huntington’s disease. ^14^ Thus, it appears that the sequence preceding a tightly folded domain impacts the proteasome’s ability to unfold the domain, with the GRR in particular directly decreasing unfolding rates of the adjacent domain, although questions remain about the mechanism.^14,20,21^

Intriguingly, a 2005 study^2^ found that a GRR sequence had different effects depending on whether degradation occurred from the N-terminus or the C-terminus of a protein. When degradation was initiated from the C-terminus, a 35-amino acid long GRR protected a folded dihydrofolate reductase (DHFR) domain if adjacent to the domain or 20 amino acids away, but not if >35 amino acids away. However, when degradation was initiated from the N-terminus, the GRR had to be 55 amino acids away from the folded DHFR domain to protect it and was unable to protect DHFR when placed closer to the domain. This discrepancy, combined with more recent structures of substrates being translocated by the proteasome, inspired us to re-examine the effects of substrate sequence composition on the processivity of degradation.

We first developed a structural model using existing cryo-EM structures that maps out the substrate translocation path for a DHFR-containing substrate (**Figure 1**). The model illustrates proteasomal features that potentially interact with the substrate as it is being translocated. Substrate residues up to 35 amino acids away from the folded DHFR domain are housed within the RP (**Figure 1A**), including, as expected, the residues that interact directly with the aromatic paddle tyrosines of the RP (**Figure 1B**). Using this model, we predict that inserting a stop signal such as a GRR within this span of residues should protect DHFR, and that this protection will be most pronounced when the stop signal is positioned such that it would interact with the aromatic paddles during unfolding of DHFR. Notably, this model predicts that, in contrast to previous results, sequences close to the domain that interact with the aromatic paddles will have the greatest effect on the proteasome’s unfolding ability, while sequences greater than 35 amino acids away will have already entered the CP when DHFR is being unfolded, so should have little effect on unfolding ability. Thus, through this study, we hope to clarify the spatial requirements for processivity and to explore additional structural features on the proteasome, apart from the aromatic paddles, which may contribute to the unfolding and translocation of substrates.

**Figure 1.**
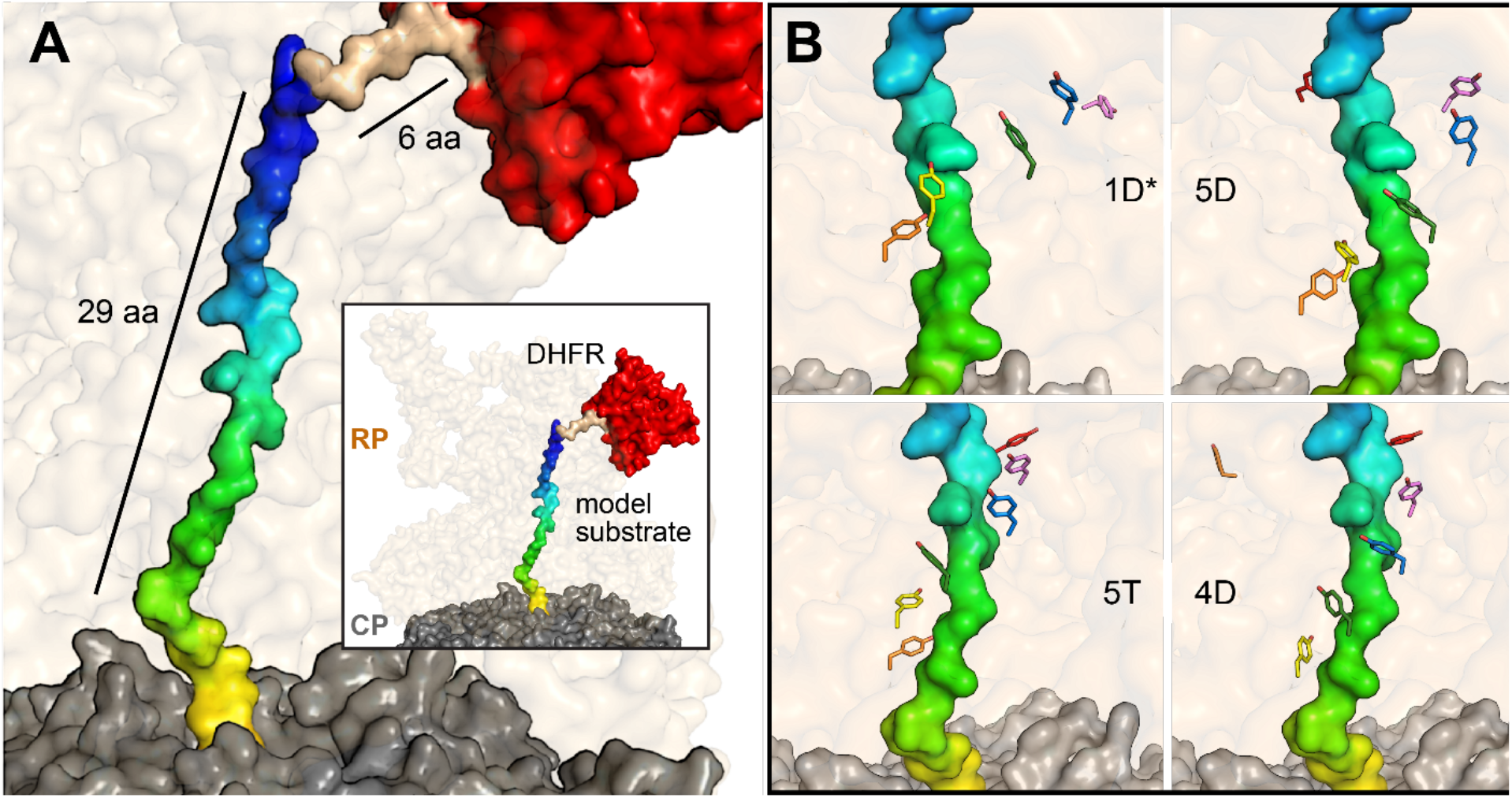
Model of substrate translocation pathway. Utilizing prior cryo-EM structures, a model of the substrate was developed to visualize the general positioning of amino acids in the translocation pathway. **A)** The CP and RP from PDB 6EF3 are shown in surface representation in gray and pale orange, respectively. The model substrate, in surface representation, is composed of the substrate from PDB 6EF3 (multicolor), along with a 6-alanine linker (brown) that models the connections between 6EF3 and the DHFR from PDB 1DRE (red), modeled at the entrance of the RP. The general positioning of the model substrate in the translocation pathway suggests that amino acids up to ∼35 amino acids away from the folded domain reside in the RP. **B)** Cryo-EM structures of the proteasome in various states were used to visualize the interactions between the aromatic paddles in those states and the substrate. The relative locations of these tyrosine residues (in sticks) that primarily interact with the amino acids from ∼16 to ∼26 away from the DHFR domain are shown. Individual tyrosines are colored the same across states as follows: Rpt1 Y283 (violet), Rpt2 Y256 (blue), Rpt6 Y222 green, Rpt3 Y246 yellow, Rpt4 Y255 orange, and Rpt5 Y255 red. 1D^*^, 5D, 5T, and 4D refer to the various states of the proteasome from PDB 6EF0, 6EF1, 6EF2, and 6EF3, respectively.^12^

## Results

### GRR regions impair unfolding

We began by conducting degradation assays on protein substrates containing a 35-residue GRR sequence to observe its effect on proteasome processivity. To establish a baseline, we used a control substrate R-N2D-ACTR-DHFR (**Figure 2A**). This substrate begins with an N-terminal unstructured degron (R-N2D) that serves as an initiation region and contains lysines to serve as ubiquitination sites (**Supplementary Table S2**). Following the R-N2D sequence, we incorporated a natively unfolded ACTR domain^22^ spanning ∼70 amino acids (which expresses well even when large portions of the sequence are changed), followed by a cys-containing linker for fluorescent labeling and then finally a difficult-to-unfold DHFR domain. The only lysines in the substrate are in the degron. The DHFR domain is stabilized by the addition of NADPH and when it is not fully degraded, the DHFR domain remains folded and is released as a stable fragment (**Figure 2A**). Indeed, the DHFR domain is completely degraded in the absence of NADPH, but some DHFR-containing fragment results from degradation in the presence of NADPH (**Supplementary Figure S2**). During the process of degradation, the proteasome binds and engages the substrate and then degrades the ACTR domain with a rate constant of *k*_deg_^full-length^ (**Figure 2A**). Upon encountering DHFR, two possible outcomes occur: 1) DHFR is unfolded, the substrate is translocated into the degradation chamber, and is degraded with a rate constant *k*_deg_^frag^, which is limited by unfolding; 2) DHFR is irreversibly released with a rate constant *k*_rel_^frag^. We use the ratio of these two outcomes (extent of complete degradation versus amount of DHFR-containing fragment formed) to determine an unfolding ability, U = k_deg_^frag^/k_rel_^frag 14^. Degradation is due to the ATP-dependent action of the proteasome, as degradation is completely prevented by the non-hydrolysable ATP analog ATPγS (**Figure 2B&C, Supplementary Figure S3**).

**Figure 2.**
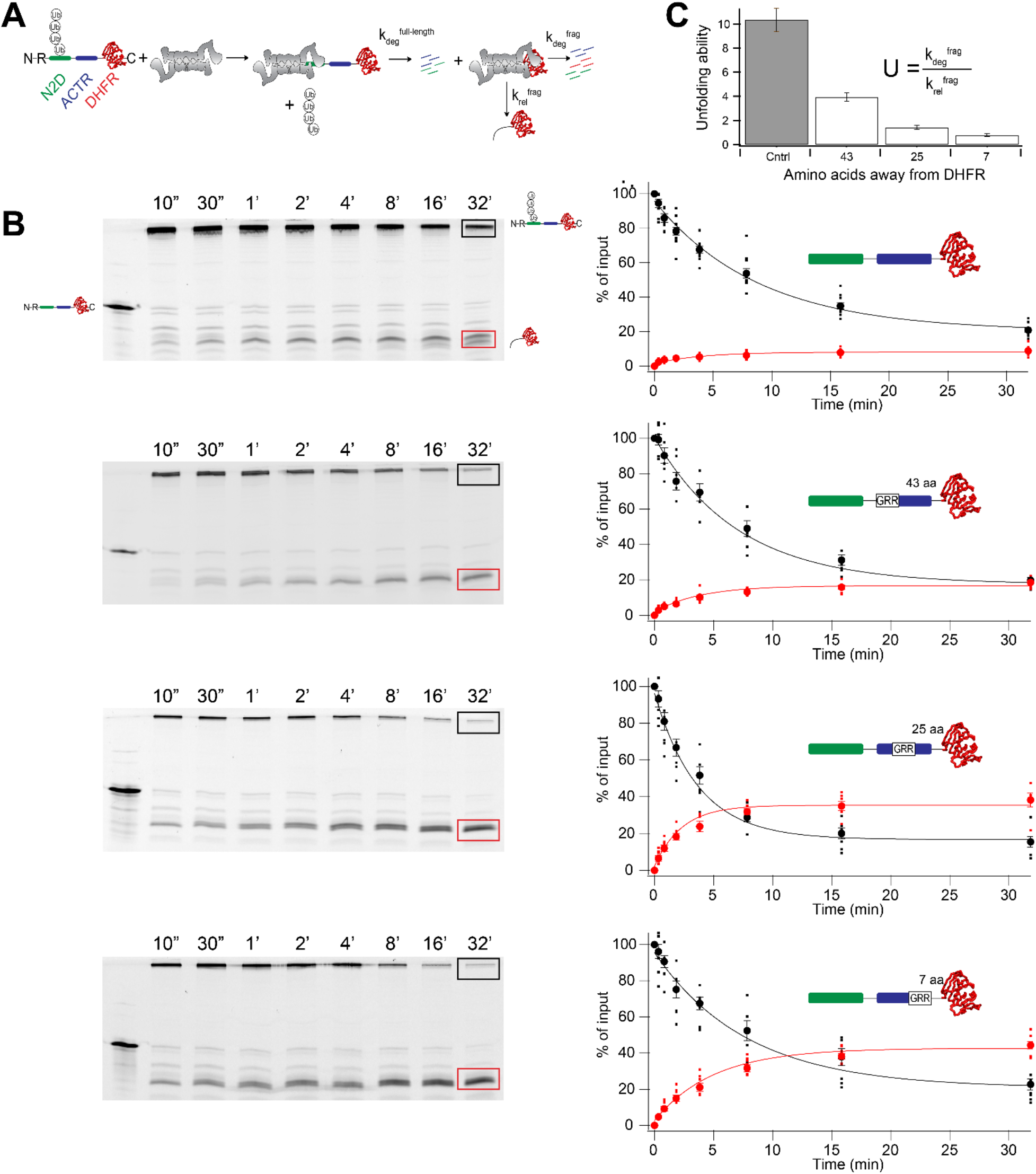
A 35-residue GRR insert impairs unfolding ability. **A)** Degradation scheme illustrating how unfolding ability is calculated. The proteasome binds ubiquitinated substrate, engaging the substrate, and degrades the ACTR domain with a rate constant of *k*_deg_^full-length^. DHFR can then either be unfolded, translocated and proteolyzed with a rate constant *k*_deg_^frag^, or released with a rate constant *k*_rel_^frag^. **B)** Representative SDS-PAGE gels (left) and quantification (right) of degradation assays. Non-ubiquitinated substrate is present in the first lane of each gel as a size reference. Disappearance of ubiquitinated substrate is shown in black, with DHFR-containing fragment in red. Individual data points are shown as dots, averages are solid symbols, error bars represent the SEM of 6 to 12 experiments, and fits are global fits to an exponential. From top to bottom: control, GRR inserted 43, 25, and 7 amino acids away from DHFR, respectively. **C)** Unfolding abilities of substrates from A. Error bars represent SEM of 6-12 experiments.

We initially generated three additional substrates by replacing sections of the ACTR domain with a GRR sequence at varying distances from the DHFR domain. The resulting substrates, ACTR_GRR8_, ACTR_GRR25_, and ACTR_GRR43_, have this GRR positioned 8, 25 and 43 residues away from DHFR. Based on our structural model, we expected the greatest reduction in unfolding ability with a substrate where the GRR is closest to DHFR (ACTR_GRR8_), while a previous study^2^ suggested that for a substrate being degraded from the N-terminus, the GRR furthest from the DHFR domain (ACTR_GRR43_) would have the greatest effect.

Insertion of the 35-residue GRR into the substrate hampered the processivity of the proteasome (**Figure 2B, C)** with all three GRR insertions resulting in >50% decrease in unfolding ability compared to the control (**Figure 2C**). The composition of the GRR likely provides fewer sidechains to facilitate the mechanical pulling force enabled by the aromatic paddles on the Rpt motors, thus reducing the unfolding ability of the proteasome by reducing the rate of unfolding of DHFR.^10,14^ In contrast to a previous report,^2^ inserting the GRR closest to DHFR caused a greater impact on unfolding ability than positioning it further away (10% of control U for ACTR_GRR8_ versus 40% for ACTR_GRR43_; p<1x10^-5^, **Supplementary Table S1**). However, in agreement with previous work ^2^, inserting a stop sequence a significant distance away from DHFR still impacts unfolding ability. Our data show that the presence of the GRR spanning 43-77 amino acids N-terminal to DHFR still diminishes processivity more than 2-fold compared to the absence of the GRR (p<1x10^-5^).

### Polyglycine regions impair unfolding differentially based on their location within the substrate

Our working hypothesis, supported by our initial GRR results, is that the region of the substrate being gripped by the proteasome during DHFR unfolding would be the most important for unfolding ability. Given that the 35-residue GRR impacted unfolding ability when positioned within a wide range of the sequence preceding DHFR, and the large size of GRR, we wished to determine if there was a narrower-grained region of importance. We therefore introduced 10 glycine (10gly) repeats (the approximate size of the region spanned by the aromatic paddles) before or into ACTR that spanned between 8-87 amino acids away from the DHFR domain. In agreement with the 35-residue GRR data, the greatest reduction in unfolding ability occurred as the 10gly regions were positioned closer to the DHFR domain (**Supplementary Figure S4 & S5, Figure 3A**). The three constructs with glycines closest to the DHFR domain, ACTR_10gly8_, ACTR_10gly13_ and ACTR_10gly18_, retained ∼20% of the unfolding ability of the control, while the other constructs were less affected (**Supplementary Table S1**). In our structural model, amino acids spanning ∼16-26 amino acids away from DHFR interact with the Rpt aromatic paddles (**Figure 3D**). Thus, the glycine stretches that have the biggest effect on unfolding map onto the regions where the aromatic paddles are predicted to interact with the substrate during unfolding, e.g., from the insert spanning from 8 to 17 aa from the DHFR through the one spanning 18 to 27 aa from DHFR.

**Figure 3.**
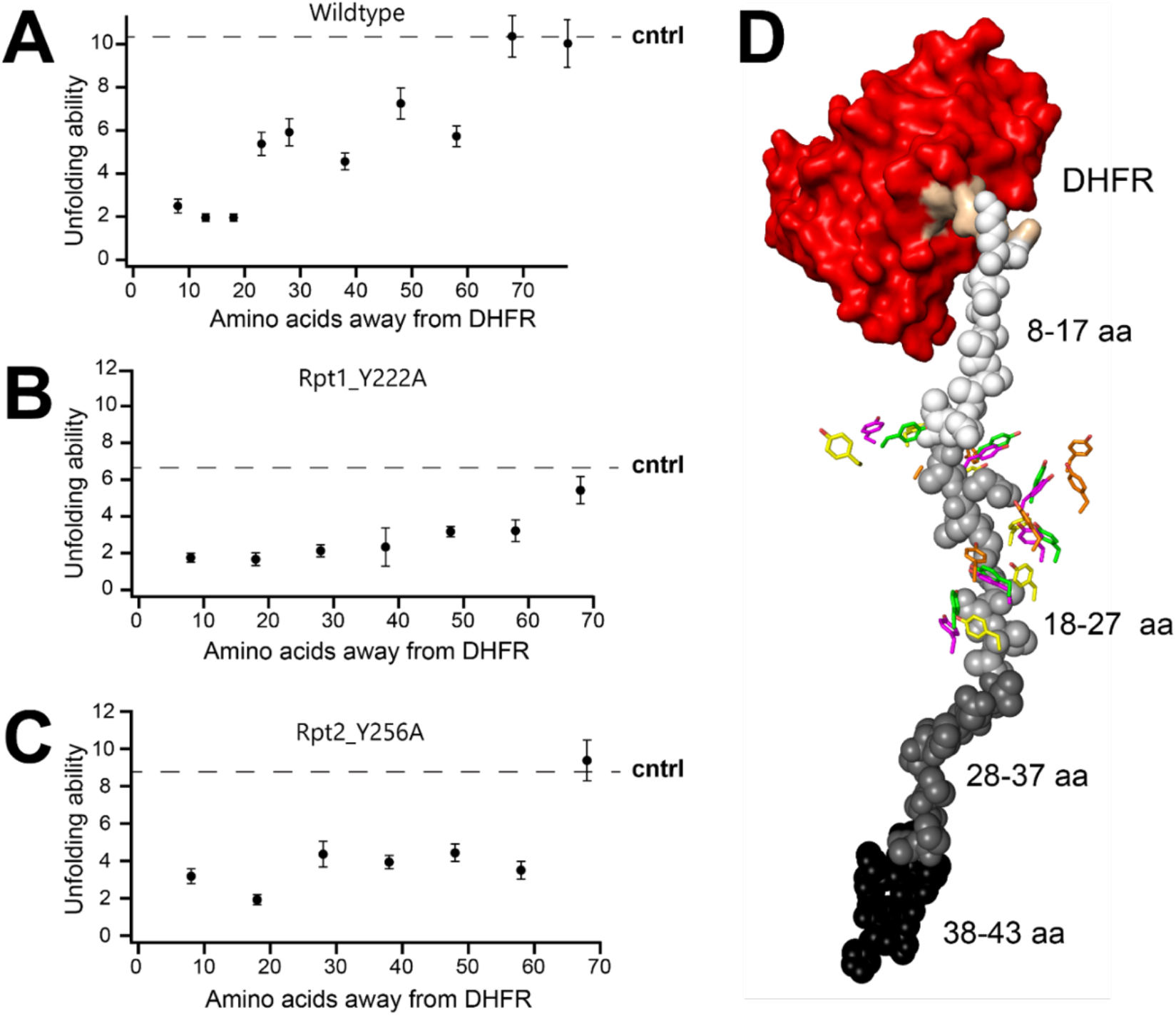
10-residue polyglycine inserts impair unfolding ability, particularly when aligned with aromatic paddles. **A)** Unfolding abilities of WT proteasome plotted against distance of inserted 10gly sequence from the DHFR domain. Dashed line is the unfolding ability of the control construct, R-N2D-ACTR-DHFR. **B)** & **C)** As in A, but with Rpt1_YA and Rpt2_YA proteasome, respectively. For A-C, error bars represent SEM of between 4 to 12 experiments. **D)** Model of the substrate with the location of polyglycine repeats, ACTR_10gly8_ - ACTR_10gly38_, mapped onto the substrate in shades of gray. Most interactions between Rpt aromatic paddles and 10gly repeats occur between 16 and 26 amino acids away from DHFR. Rpt aromatic paddles are colored according to the various states of the proteasome as follows: 1D^*^ (orange), 5D (green), 5T (magenta), and 4D (yellow).

The least processively degraded substrate, ACTR_10Gly18_, had an unfolding ability of ∼2. For this substrate, degradation is twice as fast as release of fragment, so 2/3 of the engaged substrate is degraded, and 1/3 is released as a DHFR-containing fragment. The observation that there is still substantial unfolding and degradation suggests that although the glycine stretch makes the substrate slippery, some grip is still achieved. Indeed, a highly stable domain is required to stall the proteasome even in the presence of 10gly inserts, as without NADPH to stabilize DHFR, this substrate is essentially completely degraded (**Supplementary Figure S2**).

Even though the largest effect on unfolding was observed with polyglycine insertions closer to the DHFR domain, inserting glycine tracts between 8-67 amino acids away from DHFR compromised processivity, spanning a range of impact of ∼60 amino acids. For example, ACTR_10gly58_ still resulted in a 55% reduction in unfolding ability. Polyglycine beyond 68 residues away from DHFR did not affect processivity (**Figure 3A**). The large sequence area that affects unfolding ability beyond the region 16-26 residues from DHFR suggest that other parts of the proteasome facilitate translocation of the substrate. Alternatively, it might be possible for force to be transmitted to DHFR even before it is “stuck” at the mouth of the translocation channel due to interactions with other portions of the proteasome, which could lead to different portions of the substrate interacting with the aromatic paddles during DHFR unfolding.

### Rpt aromatic paddle proteasome mutations have an altered unfolding ability profile

To distinguish between these possibilities, we assessed the processivity of the polyglycine substrates with proteasomes containing aromatic paddle mutations (Rpt1_Y222A and Rpt2_Y256A). These mutations have been shown to decrease the ability of the proteasome to unfold difficult-to-unfold proteins, but do not substantially impair degradation of less stable substrates.^10^ We reasoned that mutating a single pore-loop tyrosine to a smaller alanine (leaving five functional) have a smaller effect in the context of a glycine stretch (where grip is already weakened) than it will in the context of a “WT” more complex sequence; that is the effects of the two changes (mutation of pulling element and mutation of gripped sequence) will be sub-additive if they are working together in the same pulling step, while if they are in different locations (interfering with two pulling steps simultaneously) there will be a larger effect of mutation. A smaller effect of polygly in the region interacting with the pore loops would then lead to a “flatter” profile for paddle mutants than WT proteasome.

Indeed, we observe that the unfolding ability with substrates where the polyglycine region likely contacts the Rpt paddles (ACTR_10gly8_ and ACTR_10gly18_) are very similar between the WT and mutant proteasomes (**Figure 3B, C, Supplementary Figures S6-S9**). This result suggests that making a mutation to the aromatic paddles that interferes with pulling of the substrate has little additional effect when the polyglycine tract is introduced. In contrast, for substrates that might instead interact with other proteasomal elements, there is a pronounced difference between WT and mutant proteasomes. The mutant proteasomes result in a substantial reduction in U for these substrates (**Figure 3B, C**), flattening the profile. This effect outside the paddle-interacting region suggests that the combination of weakening the aromatic paddles as they interact with a “normal” sequence and providing a 10gly sequence that likely weakens other proteasomal interactions add together to greatly weaken the proteasome’s ability to unfold the substrate.

## Discussion

The study of how the proteasome recognizes and degrades substrates is an ongoing area of research, with implications in a variety of pathologies. Understanding the molecular mechanisms involved in the processing of substrates could enable advancements in regulating proteolysis for the treatment of these diseases.^23,24^ Here, we present evidence for some of the structural features that influence proteasome processivity by focusing primarily on the translocation pathway of the substrate as assessed using glycine-rich or poly-glycine sequences. Our results confirm that as the proteasome moves along the polypeptide sequence, its processivity is influenced by the region that comes before a tightly folded domain. This is likely due to the proteasome’s compromised grip on the substrate which hinders unfolding and subsequent degradation, leading to an increase in fragment formation. In contrast to previous findings which saw fragment formation when a GRR was inserted starting at least 55 amino acids away from DHFR we see highest fragment formation when insertions lie closer to the DHFR domain, within 8-27 amino acids away, and no processivity defect beyond 68 amino acids; it is possible that differences between our fully purified yeast proteasome assays and the *in vitro* reticulocyte lysate system used previously (which both provided ubiquitination machinery and proteasome) could explain these differences. First, we used a lysine-free DHFR in our assay, while previous work used DHFR with multiple surface lysines. Differential ubiquitination of the DHFR domain in lysate depending on the location of the GRR could affect the results if GRR near DHFR led to increased ubiquitination of DHFR and thus better degradation either via re-targeting or direct destabilization of the domain. Lysate systems also contain chaperone proteins, and differential effects of the GRR on chaperone function could also affect the observed extent of partial degradation. Differences in substrate architecture could also lead to, for example, degradation initiation from an internal site via a loop in the lysate system, which would be expected to lead to differences in grip during unfolding of DHFR. Finally, we cannot exclude the possibility that mammalian and yeast proteasome have somewhat different degradation mechanisms that lead to differential effects, although structural models of translocation are similar for mammalian and yeast proteasome. Nonetheless, our data are broadly consistent in that in both studies a GRR far from the DHFR domain can impact unfolding ability.

Collectively, these data provide a relative range of ∼70 residues preceding a folded domain as being important for pulling/unfolding of the substrate and identifies the regions that are most impactful.

These results concur with our structural model (**Figure 3D**), in which substrates containing polyglycines between 8-27 amino acids away from DHFR produced the lowest unfolding ability. One limitation of this model is that floppiness, elasticity or partial unfolding of the first several residues of DHFR could allow the 10gly of ACTR_10gly8_ to interact with more aromatic paddles than predicted. Such a mechanism might explain, for example, why ACTR_10gly8_ and ACTR_10gly18_ result in similar unfolding ability even though our model shows less overlap with the aromatic paddles for the glycines on ACTR_10gly8_ than ACTR_10gly18_. In addition, our model assumes that unfolded portions of substrates are in a similar conformation within the RP channel to those observed previously in structures; it remains possible that the substrate might take on additional conformations (i.e. loops, etc.) that could allow different portions of the substrate to contact the aromatic paddles during DHFR unfolding. Although substrate flexibility or conformational effects could explain small deviations from our model, when combined with our data with aromatic paddle mutants, it seems most likely that the smaller (but non-zero) effects of polyglycine regions further outside the paddle area are due to additional regions of the proteasome that are important for unfolding.

What could these other interactions be? Most of the interactions between the substrate and the surrounding residues (up to 5 Å away in cryo-EM structures) occur between the substrate and the aromatic paddles. The pore-2 loops located near the aromatic paddles also interact with the substrate to a smaller extent (**Supplementary Figure S10**) and have previously been proposed to interact with the substrate’s backbone.^12^ The location of both the aromatic paddles and pore-2 loops are consistent with the spatial requirements for substrate unfolding observed in our kinetic studies (**Figure 4A**). However, there are other potential interactions between the substrate and proteasome residues both before and after the Rpt aromatic paddles. In the substrate region closest to the DHFR domain, His111, Phe114, and Trp117 of Rpn11 appear to interact with the substrate (**Figure 4B**). The deubiquitinase Rpn11 sits at the entrance of the substrate channel pore, where substrate deubiquitination is coupled to degradation by the mechanical force of translocation.^25–27^ As previously reported, the Rpn11 catalytic groove is aligned with the route of the substrate pathway as it approaches the motor proteins.^12^ Therefore, it is unsurprising that these residues, many of which play catalytic roles in deubiquitination,^28^ could also be implicated in translocation, although deconvoluting their roles in deubiquitination versus translocation might prove difficult. Two or three glycines from the polyglycine tract of ACTR_10gly8_ interact at this region, and another two interact with the aromatic paddle region, which may be contributing to the low unfolding ability of this substrate.

**Figure 4.**
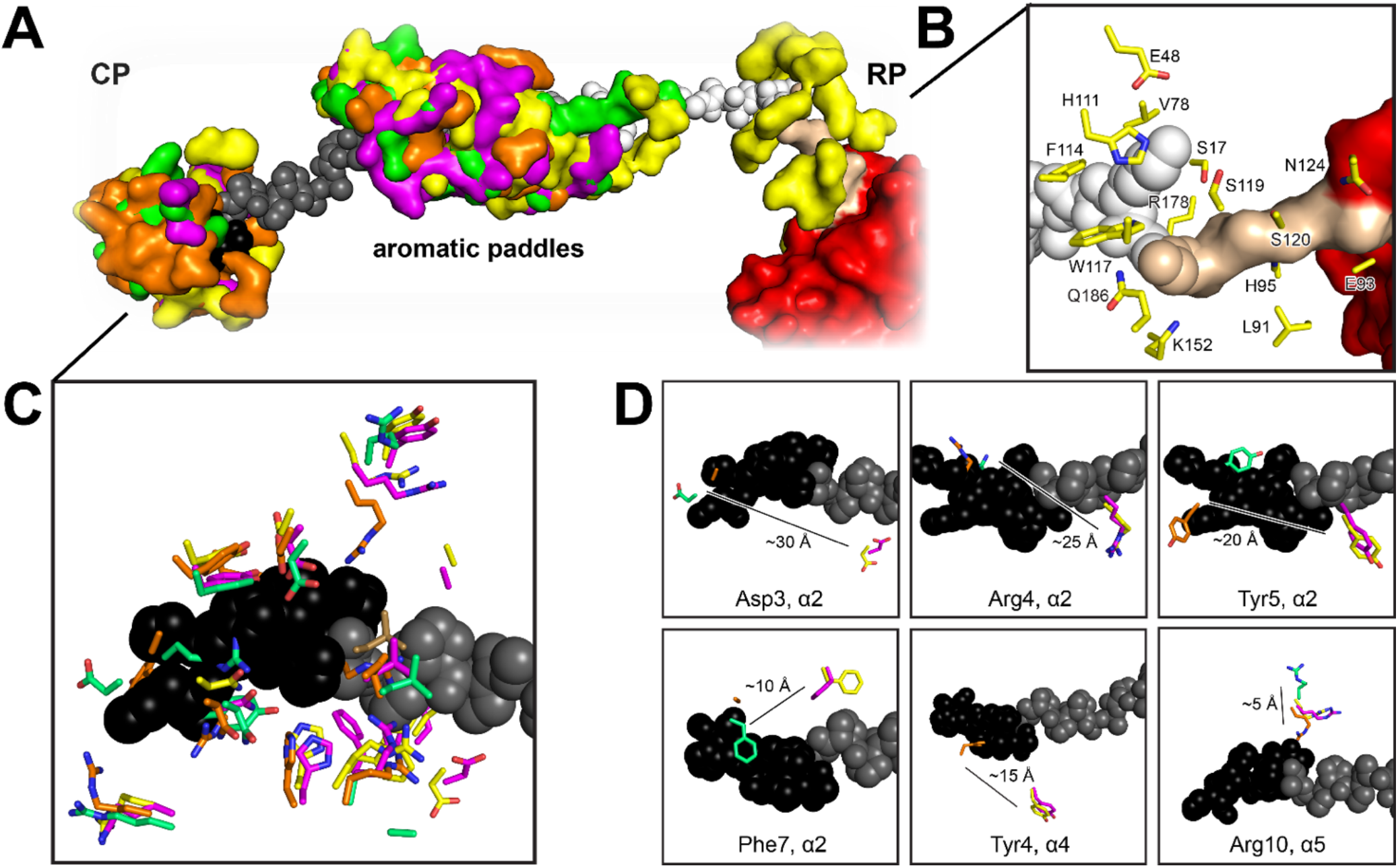
Interactions between the substrate and the surrounding residues in the translocation pathway. **A)** Residues within 5 Åof the model substrate are depicted in a surface view, with the various colors representing proteasome structures in various conformations (colored according to **Figure 3D**). Most of the interactions occur in the middle of the translocation pathway, where the Rpt aromatic paddles and the pore-2 loops are. **B)** Residues that are closer to the RP entrance than the Rpt aromatic paddles that potentially interact with the substrate. **C)** Selected residues in the CP N-terminal gating region that may interact with the substrate. **D)** Movement of residues from the alpha-2 (Asp3, Arg4, Tyr5, Phe7), alpha-4 (Tyr4), and alpha5 (Arg10) subunits of the CP between various proteasome conformations are shown, suggesting they are candidates for involvement in substrate translocation.

On the other side of the aromatic paddles, there is another cluster of potentially interacting amino acids that belong to the α subunits of the CP (**Figure 4C)**. The α subunits are located in the outer rings of the degradation chamber, and their N-termini serve as a gate, controlling the entry of substrates into the proteolytic core.^29^ Notably, Phe7 of α2 appears to move ∼10 Å as it traverses through various substrate-bound conformations providing evidence for a potential role in translocation. The observation of such a large conformational change suggests that we may be missing other amino acids that could contact the substrate in translocation conformations that have yet to be observed. One potential example is Tyr5 of α2 which appears to move ∼20 Å between various substrate-bound conformations and in one conformation (5D state), occupies the location where the substrate would be if it entered the CP (**Figure 4D**). Tyr5 and Phe7 (**Figure 4D**) were previously identified as a part of a conserved N-terminal motif implicated in substrate translocation based on *in vivo* degradation defects upon removal of the N-terminal tail in yeast.^23^ Six of the glycines in ACTR_10gly38_ map onto this region of the translocation pathway, so the observation that ACTR_10gly38_ has a significantly lower unfolding ability than ACTR_10gly28_, despite ACTR_10gly28_ being closer to the DHFR domain (p = 0.018, **Supplementary Table S1**), suggests that interactions between this region of the substrate and the α subunits are indeed important for translocation.

It is intriguing that 10gly insertions even further from the DHFR domain than the α subunits in our model are able to impact unfolding ability. Although current structures do not visualize substrate past the α subunit gating region, the active sites of the ß subunits, where the substrate is hydrolyzed, are ∼70 amino acids away from DHFR (**Supplementary Figure S11**). The small but significant effects of ACTR_10gly48_ and ACTR_10gly58_ (which extend to 67 amino acids away from DHFR) suggest that additional regions of the CP past the gating region are contributing to translocation. Following the gating region, the substrate first passes through the antechamber then through a second aperture in the β subunits to undergo hydrolysis. The antechamber provides interactions that prevent secondary structure formation so that the substrate can be accessible to the catalytic sites,^30^ but it is unclear whether these interactions contribute to translocation. Indeed, we observe that a 10gly insertion in this region (48-57 away from DHFR) has a relatively higher unfolding ability compared to the flanking 10gly insertions (**Figure 3A, Supplementary Table S1**). Next, the substrate encounters the ß subunit active sites. It has been proposed that peptide bond cleavage by the catalytic threonines can provide a pulling force sufficient to unfold some substrates.^31^ Therefore, glycine stretches within the ß subunits could interfere either with steric pulling interactions or with hydrolysis itself, and either way could reduce mechanical unfolding.^31,32^ Our observation that 10gly spanning residues 58-67 (ACTR_10gly58_) yielded similar unfolding abilities as residues spanning 23-32 (ACTR_10gly23_) and 28-37 (ACTR_10gly28_) thus indicates that sequences that map onto the β subunits likely also influence unfolding of the substrate by impairing the translocation mechanism.

Our results provide a clearer picture of the architecture within the substrate channel by highlighting the importance of the Rpt aromatic paddles and providing support for additional interactions that may be involved in translocation. When degradation begins from the N-terminus of the substrate, we show that amino acids that are ∼8-27 residues from a folded domain have the most influence on unfolding, primarily attributed to the work of the Rpt aromatic paddles. These detailed spacing requirements set the stage for future experiments in which the sequence or compositional preferences of the aromatic paddles or other possible pulling elements to be probed, and have implications for the identification of additional proteins that are partially degraded by the proteasome in cells, and for the further elucidation of proteasome-substrate interactions during unfolding and translocation.

## Materials and methods

### Substrate modeling

PDB 6EF3, which contains the CP, RP and substrate bound at the RP, was used to develop the model DHFR-containing substrate. DHFR from PDB 1DRE was initially docked at the RP substrate entry site using UDock2^33^. Since docking aims to optimize binding interactions when the proteins interface without additional forces, subsequent manual modeling of the positioning of DHFR was performed to better simulate DHFR being pulled into the channel., and a 6-residue polyalanine linker was modeled in to connect the DHFR domain and the substrate from PDB 6EF3. In the original structure (PDB 6EF3), the substrate is oriented with the C-terminus towards the CP and the N-terminus towards the RP, therefore our model has the DHFR domain grafted onto the N-terminus of the 6EF3 substrate. The coordinates of the model substrate bound to PDB 6EF3 are included in the Supplementary Information.

### Substrate Constructs

A plasmid encoding His-SUMO-R-N2D-ACTR-DHFR was derived from pCMH39 (His-SUMO-R-Neh2Dual-BarnaseΔK-C-DHFRδ5KΔC)^34^ which contains a His-SUMO tag for purification, a degron derived from Nrf2 (N2D) containing E3 binding sites and lysine ubiquitination sites, an easy-to-unfold barnase domain, a short linker containing a unique cysteine for labeling, and a hard-to-unfold DHFR domain. Barnase was replaced with the 71 amino acid ACTR domain^22^, the natively unfolded lysine-free activation domain of the p160 transcriptional co-activator for thyroid hormone and retinoid receptors, using Gibson assembly from a codon-optimized dsDNA (IDT). All lysine residues on DHFR were mutated to arginine to prevent off-target ubiquitination using oligo-directed mutagenesis.

To construct GRR-containing substrates, residues 1-35 (ACTR_GRR43_), 18-53 (ACTR_GRR25_), and 36-70 (ACTR_GRR8_) of the ACTR domain were replaced with the 35-residue GRR from p105 using Gibson Assembly such that the space between the DHFR and the GRR was 8, 25, and 43 residues, respectively. For 10gly substrates, ten amino acid segments of the ACTR domain were replaced with glycine residues as follows: ACTR_10gly68_; ACTR_10gly58_; ACTR_10gly48_; ACTR_10gly38_; ACTR_10gly28_; ACTR_10gly23_; ACTR_10gly18_; ACTR_10gly13_; ACTR_10gly8_. The number following 10gly represents the distance between the 10 glycine repeat and DHFR and is defined as the amino acid position that the glycine tract begins. For example, ACTR_10gly8_ contains the 10gly sequence beginning at the 8^th^ position away from DHFR, with 7 residues in-between. In our model substrate, these 7 residues are made up of the 6-residue polyalanine linker, and 1 residue from the substrate of PDB 6EF3. The 10gly-containing ACTR domain sequences were synthesized as dsDNA (IDT) and inserted into the BamHI and XhoI restriction sites of His-SUMO-R-Neh2Dual-ACTR-DHFR to replace the control ACTR domain. An additional mutant, ACTR_10gly78_, was made by introducing 10xglycine between the degron and ACTR using around-the-horn PCR. Sequences of all constructs are provided in **Supplementary Table S2**.

#### Protein Expression, Purification, Labeling and Ubiquitination

Protein substrates were prepared as previously described.^34^ Briefly, proteins were overexpressed in BL21 (DE3) cells using autoinduction media, purified by NiNTA chromatography and the SUMO tag was cleaved and removed by NiNTA chromatography. The single cysteine on the substrate was labeled using sulfo-cyanine 5 (Lumiprobe) and the labelled protein was then purified on a Superdex 200 column (Cytiva). Cy5-labeled substrates were then ubiquitinated using the Ubc2/Ubr1 ubiquitination system, which generates K48-linked chains attached to the degron region of the substrate as described previously^34^ using a substrate concentration of 3.0 µM. Substrates were then purified via spin size-exclusion chromatography into the degradation assay buffer. Substrates were highly ubiquitinated (single band that ran at the top of the gel, see **Supplementary Figure S1**), with an estimate of at least 15 ubiquitin per substrate.

### Proteasome purification

Proteasomes were expressed and purified using a 3x-FLAG tag on the Rpn11 subunit of the RP as previously described^34^ with the addition of a 100 mM NaCl wash prior to elution. Wildtype proteasome was from the YYS40 strain, and the Rpt1_Y22A and Rpt2_Y256A mutants contained a tyrosine to alanine mutation on the Rpt1 or Rpt2 subunits.^10^

### Degradation Assays

Degradation experiments were conducted similarly to as previously described.^4^ Assays included 25 nM proteasome incubated with 20 nM substrate at 30 °C in 50 mM Tris-Cl, 5 mM MgCl2, 5% (v/v) glycerol, 1 mM ATP, 10 mM creatine phosphate, 0.1 mg/mL creatine kinase, 1 mg/mL BSA, 0.1% Tween-20, and 1% DMSO, pH 7.5, with specific time-points quenched in SDS-PAGE sample buffer. SDS-PAGE gels were imaged on a Typhoon FLA 9500 using Cy5 fluorescence and analyzed using ImageQuant (Cytiva). For each time point, the full-length and DHFR fragment bands intensities were quantified and plotted as a percentage of the full-length substrate at the start of the reaction (10 second time point). Data were globally fit to single exponentials, and an unfolding ability was determined using equation 1:

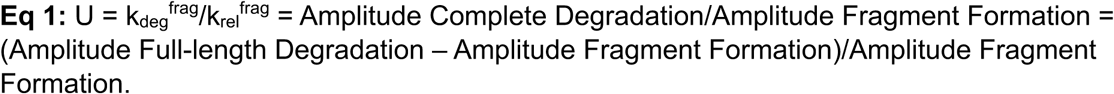

## Description of supplementary material

Supplementary data containing SDS-PAGE gels of ubiquitinations and degradation assays, degradation assay graphs, supporting figures, statistical analysis, and construct sequence information are provided.

PDB coordinates of model substrate bound to PDB 6EF3 are provided.

## Acknowledgements

The authors would like to thank Dr. Aimee Eggler, Christina M. Hurley, and members of the Kraut lab for helpful discussions and feedback. This material is based upon work supported by the National Science Foundation under Grant No. 1935596 to D.A.K.

## Conflict of Interest statement

The authors declare no conflict of interest.

## Notes

### Competing Interest Statement

The authors have declared no competing interest.

