## Supplemental Information for "Slippery sequences stall the 26S proteasome at multiple points along the translocation pathway"

**Supplementary Information**

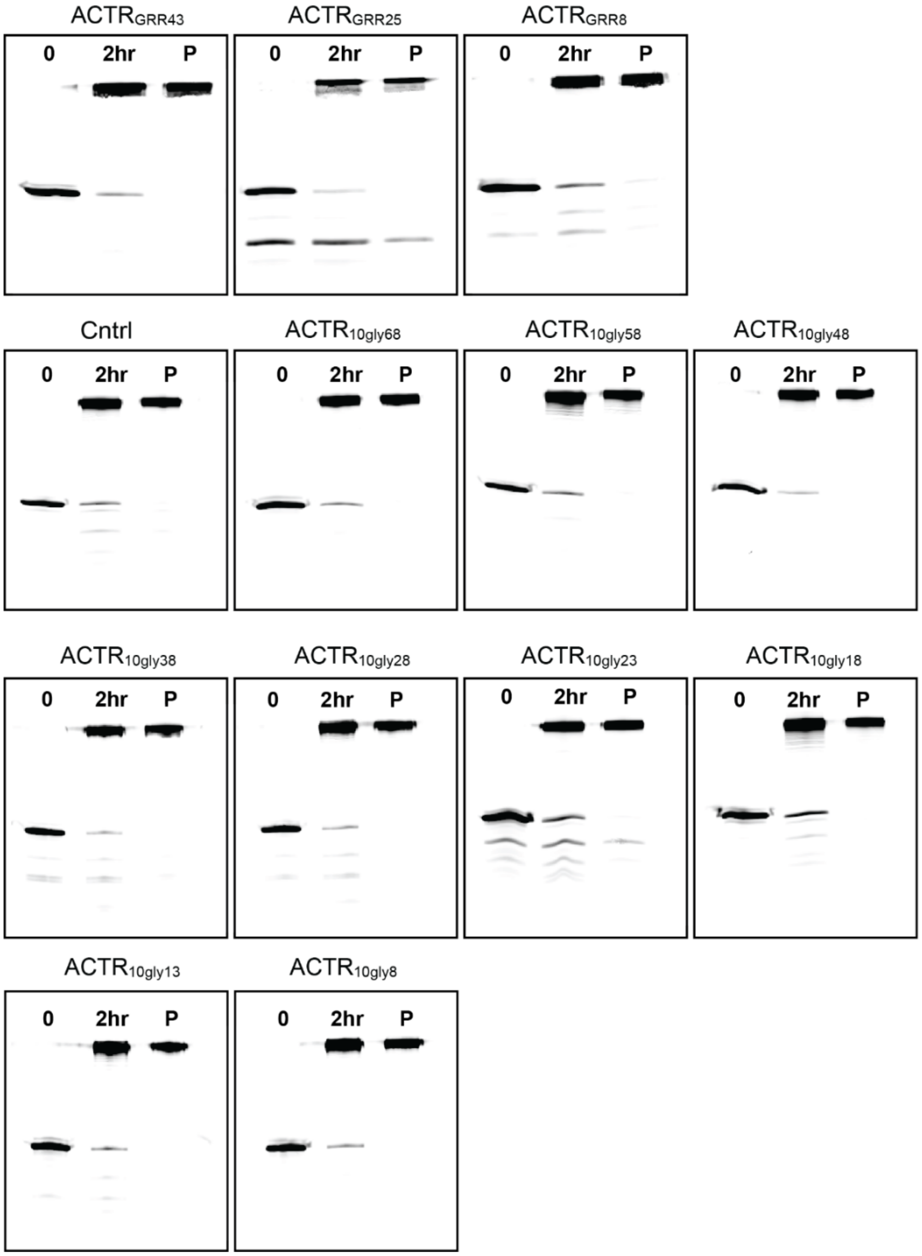

**Supplementary Figure S1. Ubiquination of substrates.** Substrates were ubiquitinated with K48-linked chains using Ubc2 (E2) and Ubr1 (E3). SDS-PAGE gels show reactions at time zero (0), after two hours (2 hr), and after purification of the ubiquitinated substrate by spin-size exclusion chromatography (P).

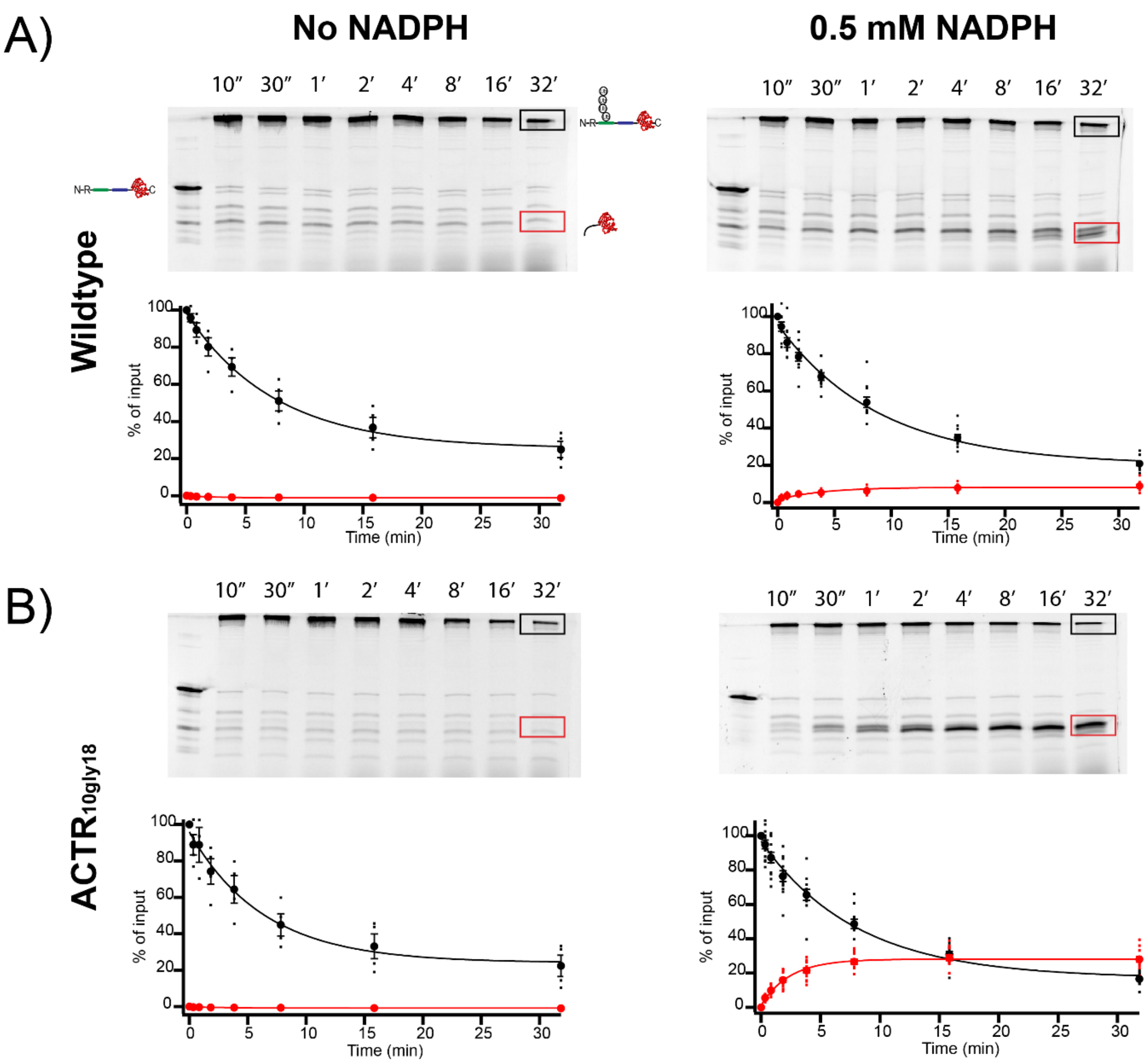

**Supplementary Figure S2. No DHFR fragment is formed when substrates are degraded in the absence of NADPH.** Experiments were conducted as outlined in **Figure 2** of the main text in the presence or absence of NADPH for **A**) the control substrate R-N2D-ACTR-DHFR and **B**) $ACTR_{10gly18}$. A representative SDS-PAGE gel is shown for each condition, with non-ubiquitinated substrate in the first lane as a size reference. Disappearance of ubiquitinated substrate is shown in black, with DHFR-containing fragment in red. Individual data points are shown as dots, averages are solid symbols, error bars represent the SEM of 3 to 12 experiments, and fits are global fits to an exponential.

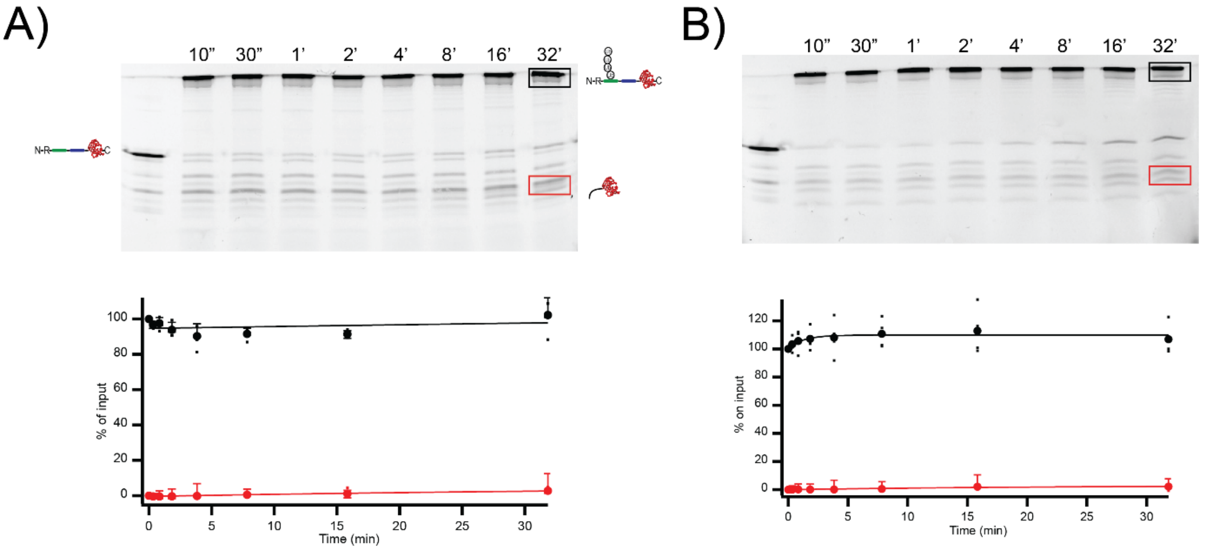

**Supplementary Figure S3. Degradation is ATP dependent**. Experiments were conducted as outlined in **Figure 2** of the main text for **A**) the control substrate R-N2D-ACTR-DHFR and **B**) $ACTR_{10gly18}$ except with 1 mM ATPγS replacing ATP and the ATP regeneration system. A representative showing that degradation is ATP-dependent. A representative SDS-PAGE gel is shown for each condition, with non-ubiquitinated substrate in the first lane as a size reference. Disappearance of ubiquitinated substrate is shown in black, with DHFR-containing fragment in red. Individual data points are shown as dots, averages are solid symbols, error bars represent the SEM of 4 experiments, and fits are global fits to an exponential.

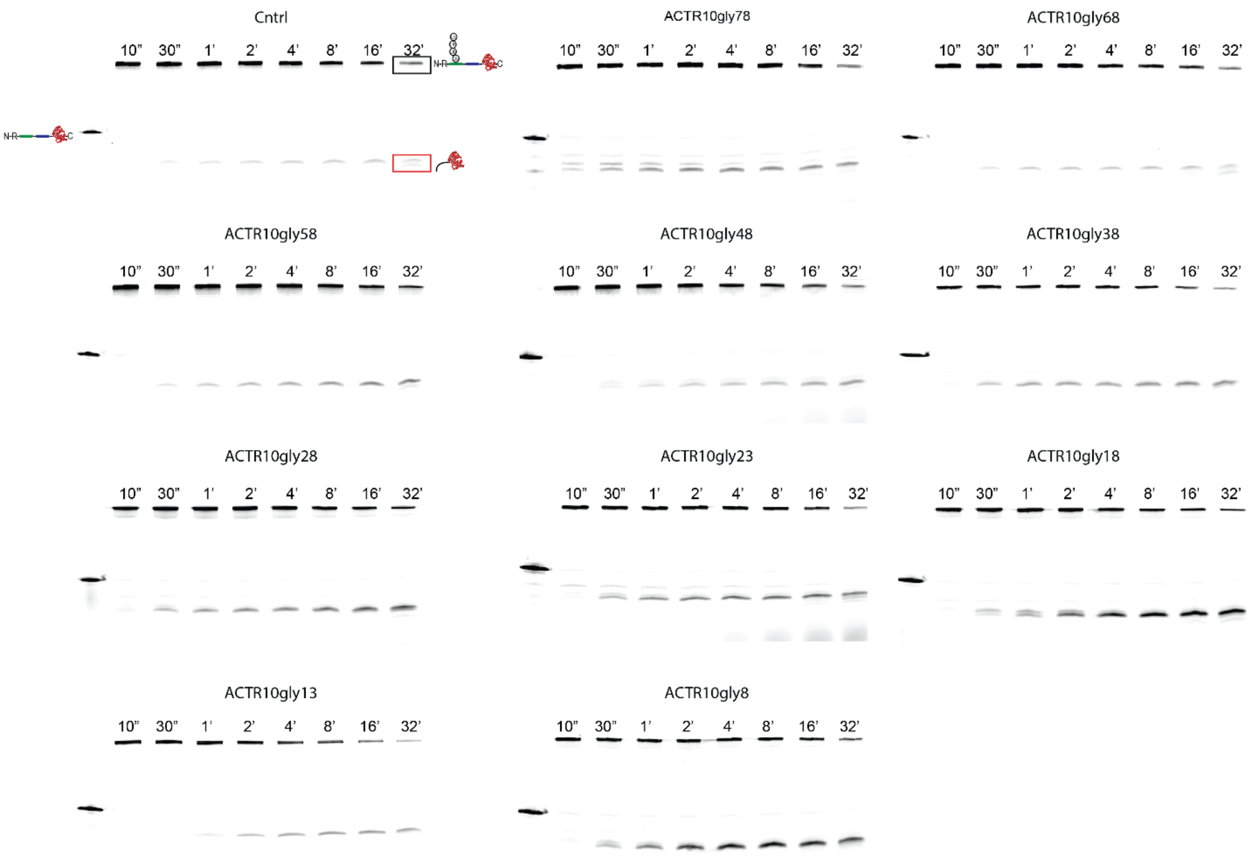

**Supplementary Figure S4. Representative gel images of degradation of polyglycine substrates by wildtype proteasome.** Experiments were conducted as outlined in **Figure 2** of the main text. Representative SDS-PAGE gels of the control substrate R-N2D-ACTR-DHFR and polyglycine substrates, with non-ubiquitinated substrate in the first lane as a size reference. Disappearance of ubiquitinated substrate is shown in black, with DHFR-containing fragment in red.

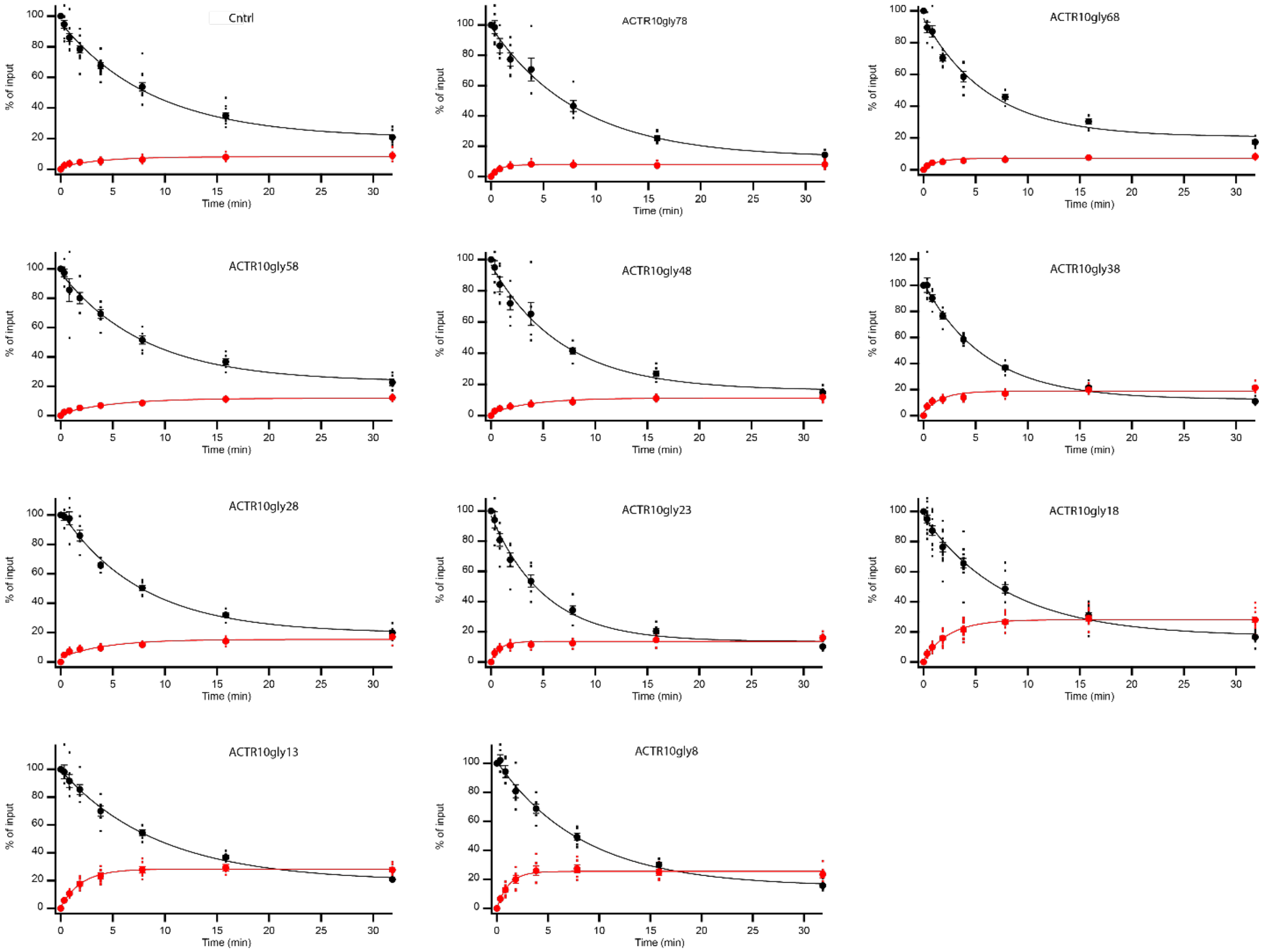

**Supplementary Figure S5. Degradation of polyglycine substrates by WT proteasome.** Experiments were conducted as outlined in **Figure 2** of the main text. Individual data points are shown as dots, averages are solid symbols, error bars represent the SEM of 4 to 12 experiments, and fits are global fits to an exponential.

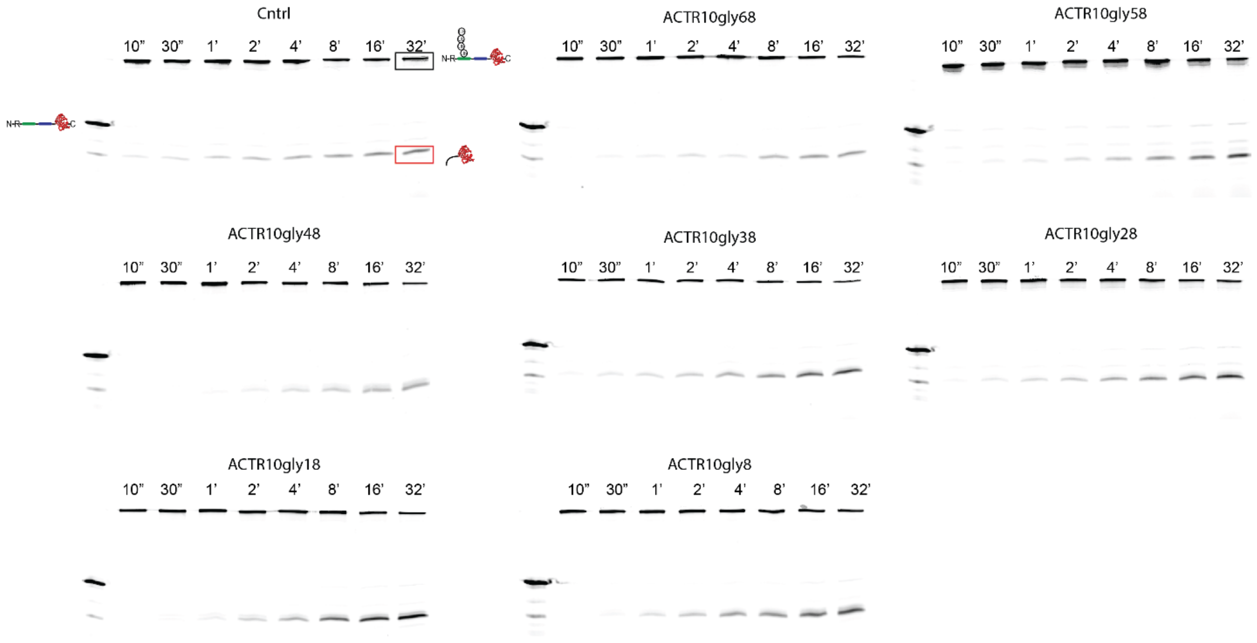

**Supplementary Figure S6. Representative gel images of degradation of polyglycine substrates by Rpt1_Y222A proteasome.** Experiments were conducted as outlined in **Figure 2** of the main text. Representative SDS-PAGE gels of the control substrate R-N2D-ACTR-DHFR and polyglycine substrates, with non-ubiquitinated substrate in the first lane as a size reference. Disappearance of ubiquitinated substrate is shown in black, with DHFR-containing fragment in red.

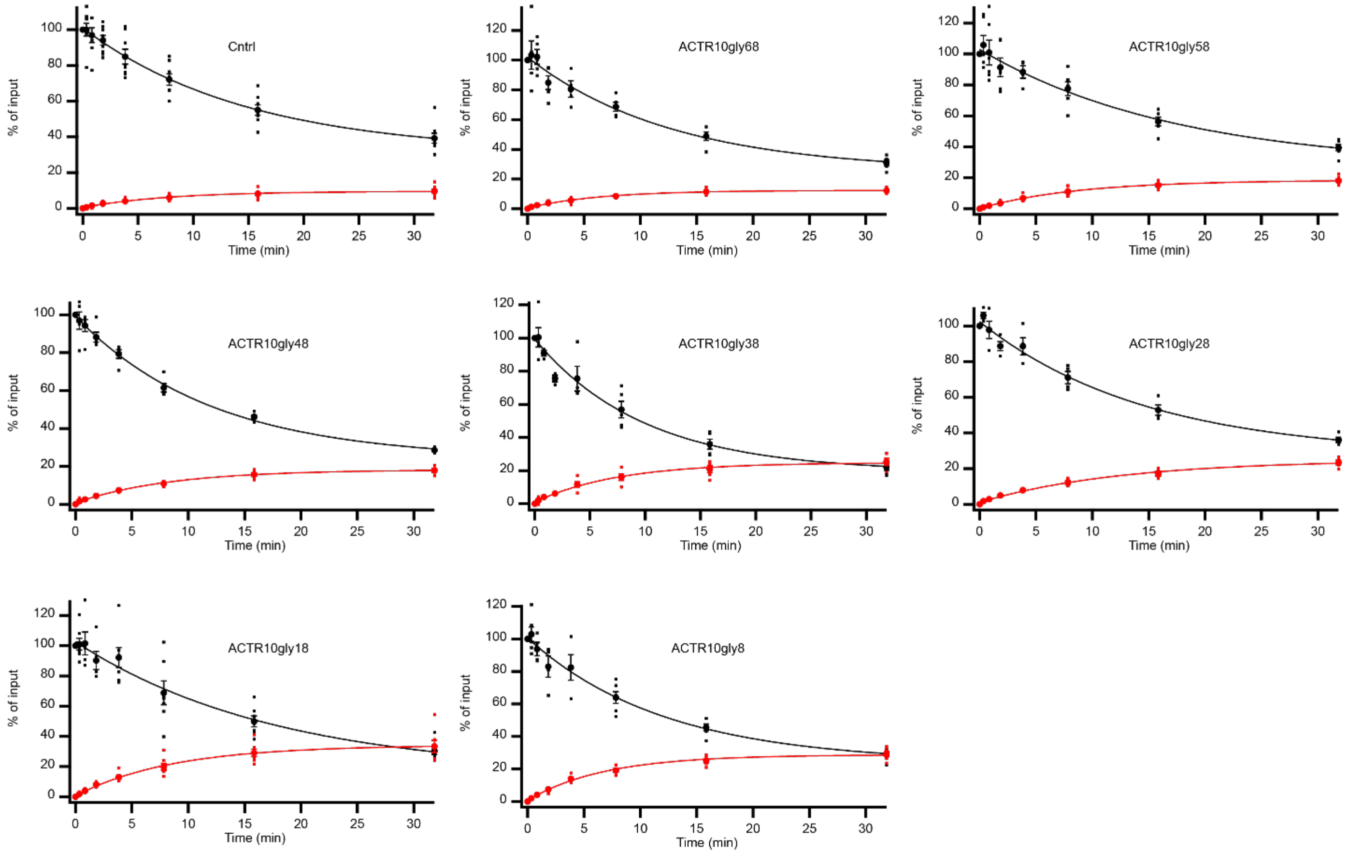

**Supplementary Figure S7. Degradation of polyglycine substrates by Rpt1_Y222A proteasome.** Experiments were conducted as outlined in **Figure 2** of the main text. Individual data points are shown as dots, averages are solid symbols, error bars represent the SEM of 3 to 8 experiments, and fits are global fits to an exponential.

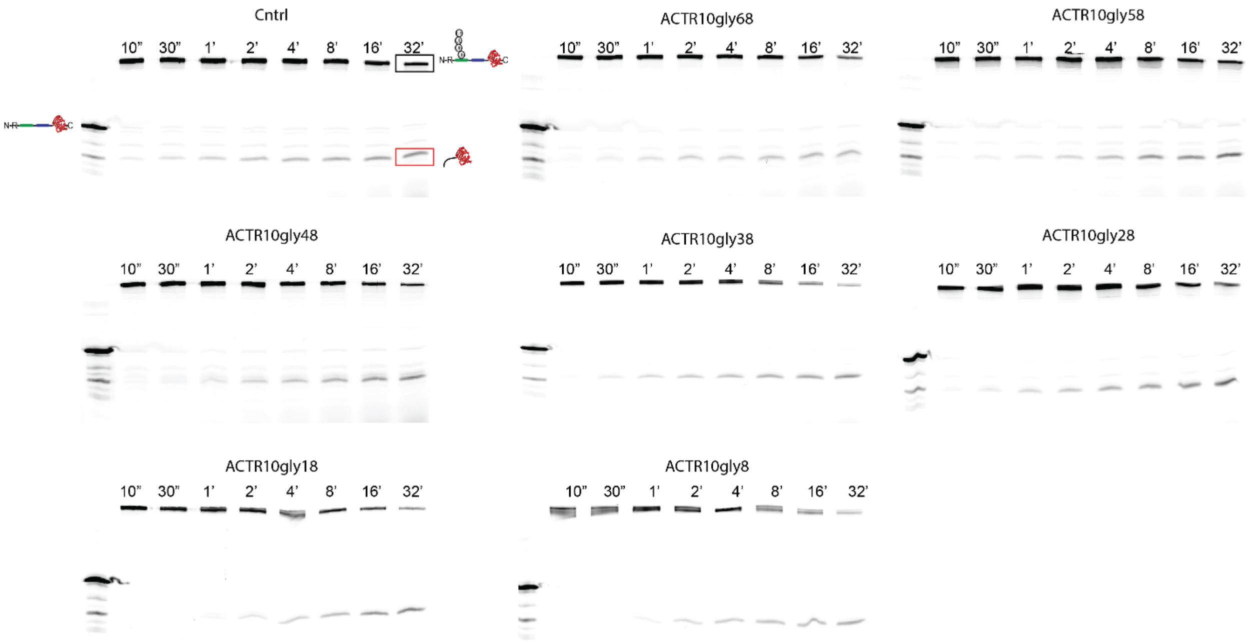

**Supplementary Figure S8. Representative gel images of degradation of polyglycine substrates by Rpt2_Y256A proteasome.** Experiments were conducted as outlined in **Figure 2** of the main text. Representative SDS-PAGE gels of the control substrate R-N2D-ACTR-DHFR and polyglycine substrates, with non-ubiquitinated substrate in the first lane as a size reference. Disappearance of ubiquitinated substrate is shown in black, with DHFR-containing fragment in red.

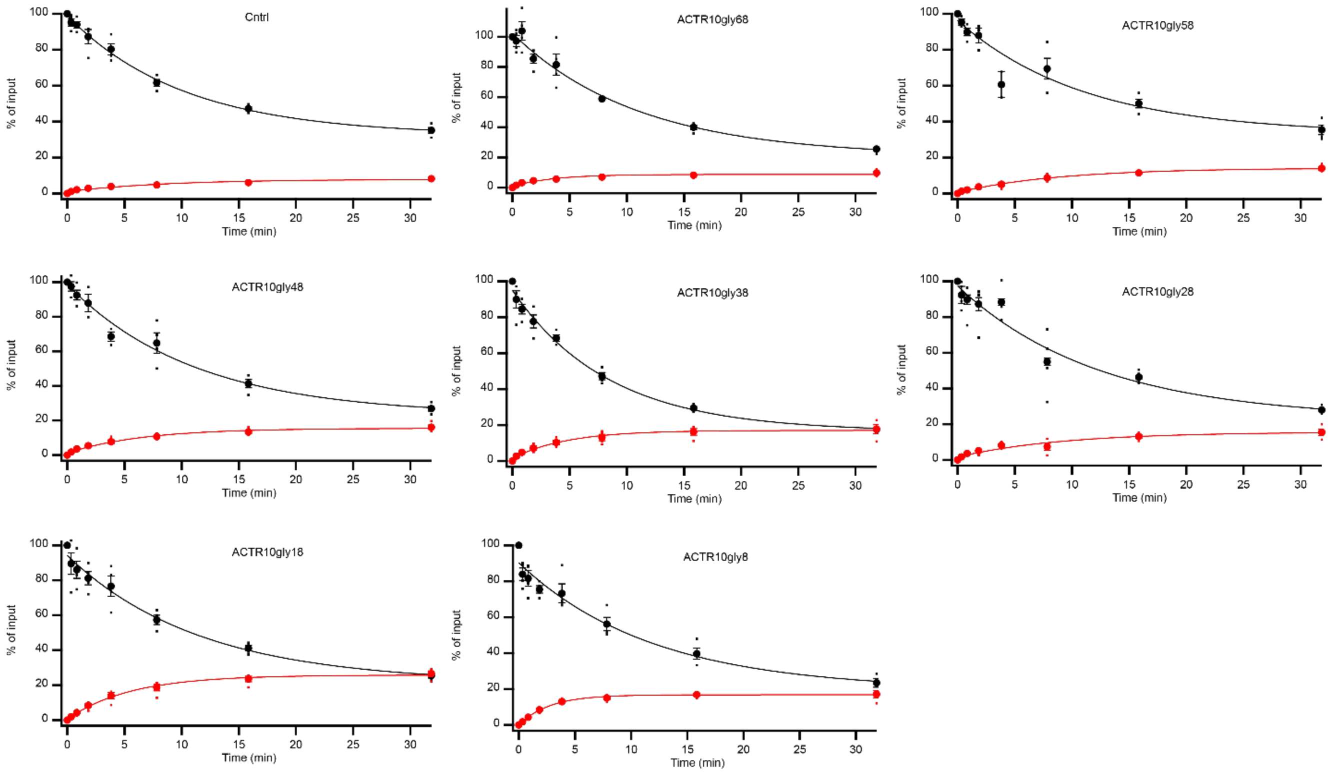

**Supplementary Figure S9. Degradation of polyglycine substrates by Rpt2_Y256A proteasome.** Experiments were conducted as outlined in **Figure 2** of the main text. Individual data points are shown as dots, averages are solid symbols, error bars represent the SEM of 3 to 8 experiments, and fits are global fits to an exponential.

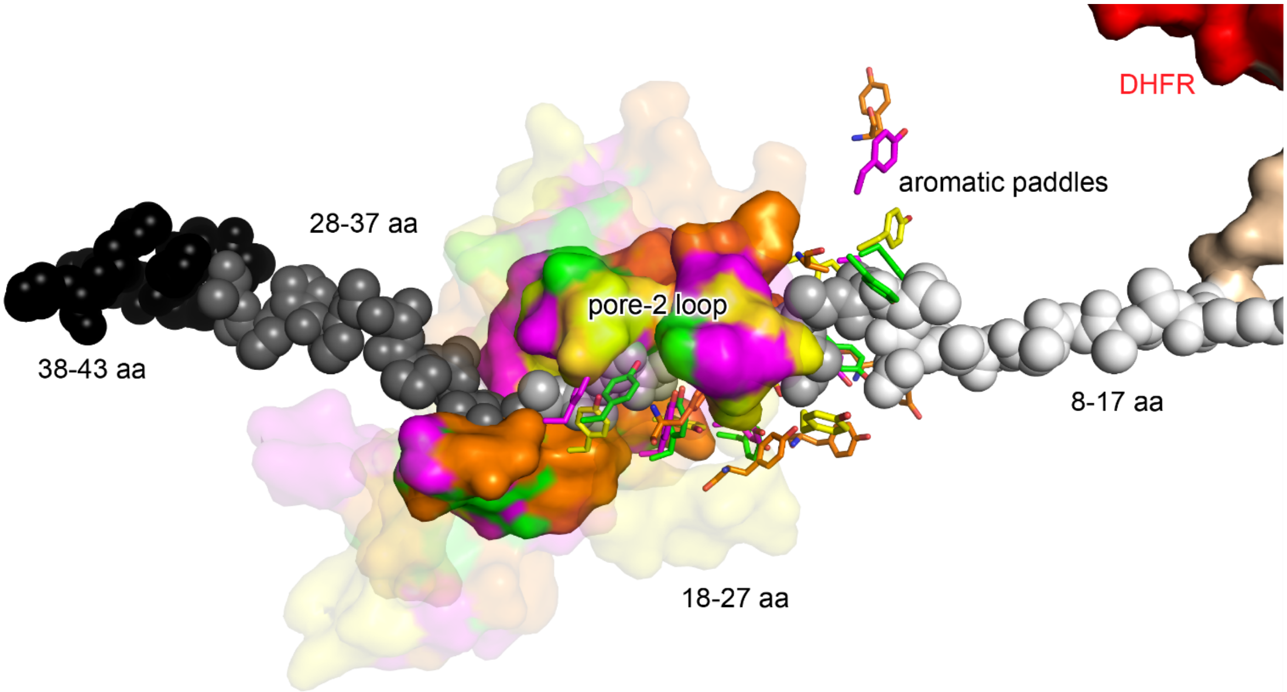

**Supplementary Figure S10. Location of the pore-2 loops in relation to aromatic paddles.** Model of the substrate with the location of 10gly repeats, $ACTR_{10gly8\text{-}10gly38}$, mapped onto the substrate in shades of gray. Rpt aromatic paddles are shown in sticks. The portion of the pore-2 loops that are within 5 Å of the substrate are shown in solid surface representations, while the rest of the pore-2 loops are shown in lighter shades. Rpt aromatic paddles and pore-2 loops are colored according to the main text **Figure 3D**.

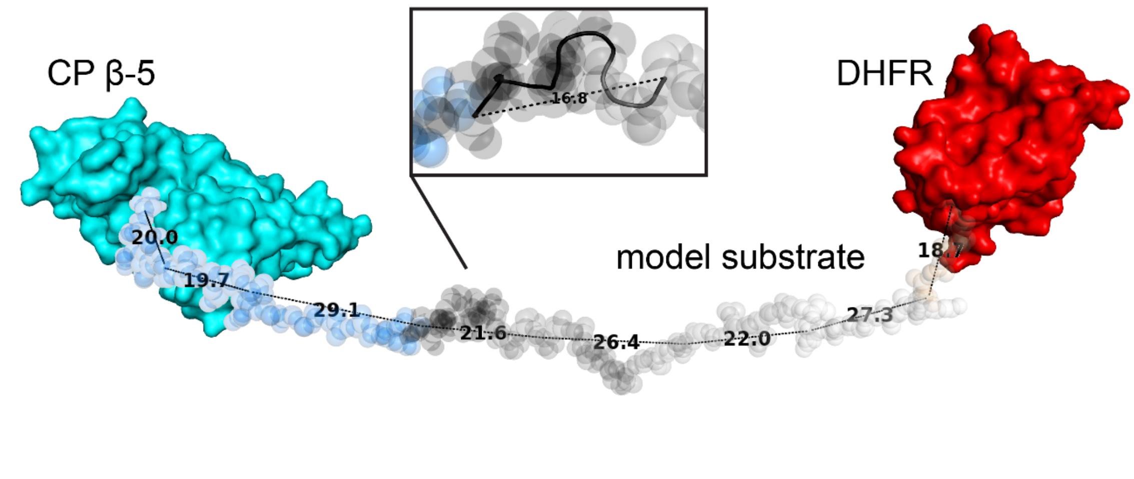

**Supplementary Figure S11. Estimated distance between DHFR and CP catalytic site.** The model substrate from **Figure 3D** was extended (blue spheres) to estimate the distance between DHFR and the catalytic threonine (Thr76) of the beta-subunit in the CP. The total distance is ~185 Å, and is covered by 70 amino acids. Additional substrate was modeled as an extended chain, with ~3 Å/amino acid, consistent with the portion of the substrate bound to the RP observed by CryoEM. Boxed area is expansion of the first 10 residues (closest to the CP) of the model substrate, showing an alpha helical secondary structure that spans 16.8 Å.

**Supplementary Table S1**. 1-way ANOVA comparison of unfolding abilities of different substrates for both WT and mutant proteasomes. A p-value below 0.05 from a Tukey-Kramer HSD post-hoc test was considered significant.

**WT proteasome, GRR substrates**

| Substrates being compared | P-value | Significance |
|---|---|---|
| ACTRGRR25 vs ACTRGRR43 | <1E-05 | ***** |
| ACTRGRR25 vs ACTRGRR8 | 1.62E-01 | |
| ACTRGRR25 vs Cntrl | <1E-05 | ***** |
| ACTRGRR43 vs ACTRGRR8 | <1E-05 | ***** |
| ACTRGRR43 vs Cntrl | <1E-05 | ***** |
| ACTRGRR8 vs Cntrl | <1E-05 | ***** |

| Annotation | p-value | Significance level |
|---|---|---|
| ***** | [0, 0.00001] | 0.00001 |
| **** | (0.00001, 0.0001] | 0.0001 |
| *** | (0.0001, 0.001] | 0.001 |
| ** | (0.001, 0.01] | 0.01 |
| * | (0.01, 0.05] | 0.05 |
| . | (0.05, 0.1] | 0.1 |
| | (0.1, 1] | 1 |

**WT proteasome, 10gly substrates**

| Substrates being compared | P-value | Significance |
|---|---|---|
| ACTR10gly13 vs ACTR10gly18 | 1.00E+00 | |
| ACTR10gly13 vs ACTR10gly23 | <1E-05 | ***** |
| ACTR10gly13 vs ACTR10gly28 | <1E-05 | ***** |
| ACTR10gly13 vs ACTR10gly38 | <1E-05 | ***** |
| ACTR10gly13 vs ACTR10gly48 | <1E-05 | ***** |
| ACTR10gly13 vs ACTR10gly58 | <1E-05 | ***** |
| ACTR10gly13 vs ACTR10gly68 | <1E-05 | ***** |
| ACTR10gly13 vs ACTR10gly78 | <1E-05 | ***** |
| ACTR10gly13 vs ACTR10gly8 | 8.88E-01 | |
| ACTR10gly13 vs | <1E-05 | ***** |
| ACTR10gly18 vs ACTR10gly23 | <1E-05 | ***** |
| ACTR10gly18 vs ACTR10gly28 | <1E-05 | ***** |
| ACTR10gly18 vs ACTR10gly38 | <1E-05 | ***** |
| ACTR10gly18 vs ACTR10gly48 | <1E-05 | ***** |
| ACTR10gly18 vs ACTR10gly58 | <1E-05 | ***** |
| ACTR10gly18 vs ACTR10gly68 | <1E-05 | ***** |
| ACTR10gly18 vs ACTR10gly78 | <1E-05 | ***** |
| ACTR10gly18 vs ACTR10gly8 | 7.40E-01 | |
| ACTR10gly18 vs Cntrl | <1E-05 | ***** |
| ACTR10gly23 vs ACTR10gly28 | 9.24E-01 | |
| ACTR10gly23 vs ACTR10gly38 | 4.91E-01 | |
| ACTR10gly23 vs ACTR10gly48 | 1.60E-04 | *** |

**Rpt1_Y222A proteasome, 10gly substrates**

| Substrates being compared | P-value | Significance |
|---|---|---|
| ACTR10gly18 vs ACTR10gly28 | 9.31E-01 | |
| ACTR10gly18 vs ACTR10gly38 | 6.11E-01 | |
| ACTR10gly18 vs ACTR10gly48 | 4.22E-03 | ** |
| ACTR10gly18 vs ACTR10gly58 | 1.47E-03 | ** |
| ACTR10gly18 vs ACTR10gly68 | <1E-05 | ***** |
| ACTR10gly18 vs ACTR10gly8 | 1.00E+00 | |
| ACTR10gly18 vs Cntrl | <1E-05 | ***** |
| ACTR10gly28 vs ACTR10gly38 | 1.00E+00 | |
| ACTR10gly28 vs ACTR10gly48 | 2.25E-01 | |
| ACTR10gly28 vs ACTR10gly58 | 1.46E-01 | |
| ACTR10gly28 vs ACTR10gly68 | <1E-05 | ***** |
| ACTR10gly28 vs ACTR10gly8 | 9.79E-01 | |
| ACTR10gly28 vs Cntrl | <1E-05 | ***** |
| ACTR10gly38 vs ACTR10gly48 | 4.06E-01 | |
| ACTR10gly38 vs ACTR10gly58 | 2.83E-01 | |
| ACTR10gly38 vs ACTR10gly68 | <1E-05 | <1E-05 |
| ACTR10gly38 vs ACTR10gly8 | 7.77E-01 | 6.77E-01 |
| ACTR10gly38 vs <1E-05 | <1E-05 | <1E-05 |
| ACTR10gly48 vs ACTR10gly58 | 1.00E+00 | 4.42E-01 |
| ACTR10gly48 vs ACTR10gly68 | 3.00E-05 | <1E-05 |
| ACTR10gly48 vs ACTR10gly8 | 1.12E-02 | 1.35E-01 |
| ACTR10gly48 vs Cntrl | <1E-05 | <1E-05 |

**Rpt2_Y256A proteasome, 10gly substrates**

| Substrates being compared | P-value | Significance |
|---|---|---|
| ACTR10gly18 vs ACTR10gly28 | 2.70E-04 | *** |
| ACTR10gly18 vs ACTR10gly38 | 2.71E-03 | ** |
| ACTR10gly18 vs ACTR10gly48 | 1.80E-04 | *** |
| ACTR10gly18 vs ACTR10gly58 | 2.83E-02 | * |
| ACTR10gly18 vs ACTR10gly68 | <1E-05 | ***** |
| ACTR10gly18 vs ACTR10gly8 | 1.30E-01 | |
| ACTR10gly18 vs Cntrl | <1E-05 | ***** |
| ACTR10gly28 vs ACTR10gly38 | 9.77E-01 | |
| ACTR10gly28 vs ACTR10gly48 | 1.00E+00 | |
| ACTR10gly28 vs ACTR10gly58 | 5.37E-01 | |
| ACTR10gly28 vs ACTR10gly68 | <1E-05 | ***** |
| ACTR10gly28 vs ACTR10gly8 | 1.82E-01 | |
| ACTR10gly28 vs Cntrl | <1E-05 | ***** |
| ACTR10gly38 vs ACTR10gly48 | 9.49E-01 | |
| ACTR10gly38 vs ACTR10gly58 | 9.71E-01 | |
| ACTR10gly38 vs ACTR10gly68 | <1E-05 | ***** |
| ACTR10gly38 vs ACTR10gly8 | 6.77E-01 | |
| ACTR10gly38 vs Cntrl | <1E-05 | ***** |
| ACTR10gly48 vs ACTR10gly58 | 4.42E-01 | |
| ACTR10gly48 vs ACTR10gly68 | <1E-05 | ***** |
| ACTR10gly48 vs ACTR10gly8 | 1.35E-01 | |
| ACTR10gly48 vs Cntrl | <1E-05 | ***** |

**WT proteasome, GRR substrates**

| Substrates being compared | P-value | Significance |
|---|---|---|

| Annotation | p-value | Significance level |
|---|---|---|
| ***** | [0, 0.00001] | 0.00001 |
| **** | (0.00001, 0.0001] | 0.0001 |
| *** | (0.0001, 0.001] | 0.001 |
| ** | (0.001, 0.01] | 0.01 |
| * | (0.01, 0.05] | 0.05 |
| . | (0.05, 0.1] | 0.1 |
| | (0.1, 1] | 1 |

**WT proteasome, 10gly substrates**

| Substrates being compared | P-value | Significance |
|---|---|---|
| ACTR10gly23 vs ACTR10gly58 | 9.97E-01 | |
| ACTR10gly23 vs ACTR10gly68 | <1E-05 | ***** |
| ACTR10gly23 vs ACTR10gly78 | <1E-05 | ***** |
| ACTR10gly23 vs ACTR10gly8 | <1E-05 | ***** |
| ACTR10gly23 vs Cntrl | <1E-05 | ***** |
| ACTR10gly28 vs ACTR10gly38 | 1.77E-02 | * |
| ACTR10gly28 vs ACTR10gly48 | 2.25E-02 | * |
| ACTR10gly28 vs ACTR10gly58 | 1.00E+00 | |
| ACTR10gly28 vs ACTR10gly68 | <1E-05 | ***** |
| ACTR10gly28 vs ACTR10gly78 | <1E-05 | ***** |
| ACTR10gly28 vs ACTR10gly8 | <1E-05 | ***** |
| ACTR10gly28 vs Cntrl | <1E-05 | ***** |
| ACTR10gly38 vs ACTR10gly48 | <1E-05 | ***** |
| ACTR10gly38 vs ACTR10gly58 | 7.38E-02 | . |
| ACTR10gly38 vs ACTR10gly68 | <1E-05 | ***** |
| ACTR10gly38 vs ACTR10gly78 | <1E-05 | ***** |
| ACTR10gly38 vs ACTR10gly8 | <1E-05 | ***** |
| ACTR10gly38 vs Cntrl | <1E-05 | ***** |
| ACTR10gly48 vs ACTR10gly58 | 4.52E-03 | ** |

**Rpt1_Y222A proteasome, 10gly substrates**

| Substrates being compared | P-value | Significance |
|---|---|---|
| ACTR10gly58 vs ACTR10gly68 | 2.00E-05 | **** |
| ACTR10gly58 vs ACTR10gly8 | 4.53E-03 | ** |
| ACTR10gly58 vs Cntrl | <1E-05 | ***** |
| ACTR10gly68 vs ACTR10gly8 | <1E-05 | ***** |
| ACTR10gly68 vs Cntrl | 3.32E-02 | * |
| ACTR10gly8 vs Cntrl | <1E-05 | ***** |

**Rpt2_Y256A proteasome, 10gly substrates**

| Substrates being compared | P-value | Significance |
|---|---|---|
| ACTR10gly58 vs ACTR10gly68 | <1E-05 | ***** |
| ACTR10gly58 vs ACTR10gly8 | 9.95E-01 | |
| ACTR10gly58 vs Cntrl | <1E-05 | ***** |
| ACTR10gly68 vs ACTR10gly8 | <1E-05 | ***** |
| ACTR10gly68 vs Cntrl | 6.07E-01 | |
| ACTR10gly8 vs Cntrl | <1E-05 | ***** |

**Supplementary Table S2.** Substrate sequence composition.

| Construct | Sequence |
|---|---|
| Neh2Dual-ACTR-C-DHFR | RDLELPPPYLPSQQDMDLIDILWRQDIDLGVSREVFDFSQRRKEYELEKQKKLEKER<br>QEQLQKEQEKAFFAQLQLDEETGEFLPIQPAQHTQSETSGSEQVSHGTQNRPLLRNS<br>LDDLVGPPSNLEGQSDERALLDQLHTLLSNTDATGLEEIDRALGIPELVNQGQALTG<br>CHLEMISLIAALAVDRVIGMENAMPWNLPADLAWFRRNTLNRPVIMGRHTWESIGRP<br>LPGRRNIILSSQPGTDDRVTWVRSVDEAIAAAGDVPEIMVIGGGRVYEQFLPRAQRL<br>YLTHIDAEVEGDTHFPDYEPDDWESVFSEFHDADAQNSHSYSFEILERR |
| Neh2Dual-ACTR$_{GRR8}$-C-DHFR | RDLELPPPYLPSQQDMDLIDILWRQDIDLGVSREVFDFSQRRKEYELEKQKKLEKER<br>QEQLQKEQEKAFFAQLQLDEETGEFLPIQPAQHTQSETSGSEQVSHGTQNRPLLRNS<br>LDDLVGPPSNLEGQSDERALNFSDSFGGGSGAGAGGGGMFGSGGGGGGTGSTGPGTG<br>CHLEMISLIAALAVDRVIGMENAMPWNLPADLAWFRRNTLNRPVIMGRHTWESIGRP<br>LPGRRNIILSSQPGTDDRVTWVRSVDEAIAAAGDVPEIMVIGGGRVYEQFLPRAQRL<br>YLTHIDAEVEGDTHFPDYEPDDWESVFSEFHDADAQNSHSYSFEILERR |
| Neh2Dual-ACTR$_{GRR25}$-C-DHFR | RDLELPPPYLPSQQDMDLIDILWRQDIDLGVSREVFDFSQRRKEYELEKQKKLEKER<br>QEQLQKEQEKAFFAQLQLDEETGEFLPIQPAQHTQSETSGSEQVSHGTQNRPLLRNS<br>LNFSDSFGGGSGAGAGGGGMFGSGGGGGGTGSTGPGEEIDRALGIPELVNQGQALTG<br>CHLEMISLIAALAVDRVIGMENAMPWNLPADLAWFRRNTLNRPVIMGRHTWESIGRP<br>LPGRRNIILSSQPGTDDRVTWVRSVDEAIAAAGDVPEIMVIGGGRVYEQFLPRAQRL<br>YLTHIDAEVEGDTHFPDYEPDDWESVFSEFHDADAQNSHSYSFEILERR |
| Neh2Dual-ACTR$_{GRR43}$-C-DHFR | RDLELPPPYLPSQQDMDLIDILWRQDIDLGVSREVFDFSQRRKEYELEKQKKLEKER<br>QEQLQKEQEKAFFAQLQLDEETGEFLPIQPAQHTQSETSGSNFSDSFGGGSGAGAGG<br>GGMFGSGGGGGGTGSTGPGLLDQLHTLLSNTDATGLEEIDRALGIPELVNQGQALTG<br>CHLEMISLIAALAVDRVIGMENAMPWNLPADLAWFRRNTLNRPVIMGRHTWESIGRP<br>LPGRRNIILSSQPGTDDRVTWVRSVDEAIAAAGDVPEIMVIGGGRVYEQFLPRAQRL<br>YLTHIDAEVEGDTHFPDYEPDDWESVFSEFHDADAQNSHSYSFEILERR |
| Neh2Dual-ACTR$_{10gly8}$-C-DHFR | RDLELPPPYLPSQQDMDLIDILWRQDIDLGVSREVFDFSQRRKEYELEKQKKLEKER<br>QEQLQKEQEKAFFAQLQLDEETGEFLPIQPAQHTQSETSGSEQVSHGTQNRPLLRNS<br>LDDLVGPPSNLEGQSDERALLDQLHTLLSNTDATGLEEIDRALGGGGGGGGGGGLTG<br>CHLEMISLIAALAVDRVIGMENAMPWNLPADLAWFRRNTLNRPVIMGRHTWESIGRP<br>LPGRRNIILSSQPGTDDRVTWVRSVDEAIAAAGDVPEIMVIGGGRVYEQFLPRAQRL<br>YLTHIDAEVEGDTHFPDYEPDDWESVFSEFHDADAQNSHSYSFEILERR |
| Neh2Dual-ACTR$_{10gly13}$-C-DHFR | RDLELPPPYLPSQQDMDLIDILWRQDIDLGVSREVFDFSQRRKEYELEKQKKLEKER<br>QEQLQKEQEKAFFAQLQLDEETGEFLPIQPAQHTQSETSGSEQVSHGTQNRPLLRNS<br>LDDLVGPPSNLEGQSDERALLDQLHTLLSNTDATGLEEIGGGGGGGGGGNQGQALTG<br>CHLEMISLIAALAVDRVIGMENAMPWNLPADLAWFRRNTLNRPVIMGRHTWESIGRP<br>LPGRRNIILSSQPGTDDRVTWVRSVDEAIAAAGDVPEIMVIGGGRVYEQFLPRAQRL<br>YLTHIDAEVEGDTHFPDYEPDDWESVFSEFHDADAQNSHSYSFEILERR |
| Neh2Dual-ACTR$_{10gly18}$-C-DHFR | RDLELPPPYLPSQQDMDLIDILWRQDIDLGVSREVFDFSQRRKEYELEKQKKLEKER<br>QEQLQKEQEKAFFAQLQLDEETGEFLPIQPAQHTQSETSGSEQVSHGTQNRPLLRNS<br>LDDLVGPPSNLEGQSDERALLDQLHTLLSNTDATGGGGGGGGGGIPELVNQGQALTG<br>CHLEMISLIAALAVDRVIGMENAMPWNLPADLAWFRRNTLNRPVIMGRHTWESIGRP<br>LPGRRNIILSSQPGTDDRVTWVRSVDEAIAAAGDVPEIMVIGGGRVYEQFLPRAQRL<br>YLTHIDAEVEGDTHFPDYEPDDWESVFSEFHDADAQNSHSYSFEILERR |

| Construct | Sequence |
|---|---|
| Neh2Dual-ACTR$_{10gly23}$-C-DHFR | RDLELPPPYLPSQQDMDLIDILWRQDIDLGVSREVFDFSQRRKEYELEKQKKLEKER<br>QEQLQKEQEKAFFAQLQLDEETGEFLPIQPAQHTQSETSGSEQVSHGTQNRPLLRNS<br>LDDLVGPPSNLEGQSDERALLDQLHTLLSGGGGGGGGGGDRALGIPELVNQGQALTG<br>CHLEMISLIAALAVDRVIGMENAMPWNLPADLAWFRRNTLNRPVIMGRHTWESIGRP<br>LPGRRNIILSSQPGTDDRVTWVRSVDEAIAAAGDVPEIMVIGGGRVYEQFLPRAQRL<br>YLTHIDAEVEGDTHFPDYEPDDWESVFSEFHDADAQNSHSYSFEILERR |
| Neh2Dual-ACTR$_{10gly28}$-C-DHFR | RDLELPPPYLPSQQDMDLIDILWRQDIDLGVSREVFDFSQRRKEYELEKQKKLEKER<br>QEQLQKEQEKAFFAQLQLDEETGEFLPIQPAQHTQSETSGSEQVSHGTQNRPLLRNS<br>LDDLVGPPSNLEGQSDERALLDQLGGGGGGGGGGGLEEIDRALGIPELVNQGQALTG<br>CHLEMISLIAALAVDRVIGMENAMPWNLPADLAWFRRNTLNRPVIMGRHTWESIGRP<br>LPGRRNIILSSQPGTDDRVTWVRSVDEAIAAAGDVPEIMVIGGGRVYEQFLPRAQRL<br>YLTHIDAEVEGDTHFPDYEPDDWESVFSEFHDADAQNSHSYSFEILERR |
| Neh2Dual-ACTR$_{10gly38}$-C-DHFR | RDLELPPPYLPSQQDMDLIDILWRQDIDLGVSREVFDFSQRRKEYELEKQKKLEKER<br>QEQLQKEQEKAFFAQLQLDEETGEFLPIQPAQHTQSETSGSEQVSHGTQNRPLLRNS<br>LDDLVGPPSNLEGQGGGGGGGGGGHTLLSNTDATGLEEIDRALGIPELVNQGQALTG<br>CHLEMISLIAALAVDRVIGMENAMPWNLPADLAWFRRNTLNRPVIMGRHTWESIGRP<br>LPGRRNIILSSQPGTDDRVTWVRSVDEAIAAAGDVPEIMVIGGGRVYEQFLPRAQRL<br>YLTHIDAEVEGDTHFPDYEPDDWESVFSEFHDADAQNSHSYSFEILERR |
| Neh2Dual-ACTR$_{10gly48}$-C-DHFR | RDLELPPPYLPSQQDMDLIDILWRQDIDLGVSREVFDFSQRRKEYELEKQKKLEKER<br>QEQLQKEQEKAFFAQLQLDEETGEFLPIQPAQHTQSETSGSEQVSHGTQNRPLLRNS<br>LDDLGGGGGGGGGGSDERALLDQLHTLLSNTDATGLEEIDRALGIPELVNQGQALTG<br>CHLEMISLIAALAVDRVIGMENAMPWNLPADLAWFRRNTLNRPVIMGRHTWESIGRP<br>LPGRRNIILSSQPGTDDRVTWVRSVDEAIAAAGDVPEIMVIGGGRVYEQFLPRAQRL<br>YLTHIDAEVEGDTHFPDYEPDDWESVFSEFHDADAQNSHSYSFEILERR |
| Neh2Dual-ACTR$_{10gly58}$-C-DHFR | RDLELPPPYLPSQQDMDLIDILWRQDIDLGVSREVFDFSQRRKEYELEKQKKLEKER<br>QEQLQKEQEKAFFAQLQLDEETGEFLPIQPAQHTQSETSGSEQVSHGTQNRGGGGGG<br>GGGGVGPPSNLEGQSDERALLDQLHTLLSNTDATGLEEIDRALGIPELVNQGQALTG<br>CHLEMISLIAALAVDRVIGMENAMPWNLPADLAWFRRNTLNRPVIMGRHTWESIGRP<br>LPGRRNIILSSQPGTDDRVTWVRSVDEAIAAAGDVPEIMVIGGGRVYEQFLPRAQRL<br>YLTHIDAEVEGDTHFPDYEPDDWESVFSEFHDADAQNSHSYSFEILERR |
| Neh2Dual-ACTR$_{10gly68}$-C-DHFR | RDLELPPPYLPSQQDMDLIDILWRQDIDLGVSREVFDFSQRRKEYELEKQKKLEKER<br>QEQLQKEQEKAFFAQLQLDEETGEFLPIQPAQHTQSETSGSGGGGGGGGGGPLLRNS<br>LDDLVGPPSNLEGQSDERALLDQLHTLLSNTDATGLEEIDRALGIPELVNQGQALTG<br>CHLEMISLIAALAVDRVIGMENAMPWNLPADLAWFRRNTLNRPVIMGRHTWESIGRP<br>LPGRRNIILSSQPGTDDRVTWVRSVDEAIAAAGDVPEIMVIGGGRVYEQFLPRAQRL<br>YLTHIDAEVEGDTHFPDYEPDDWESVFSEFHDADAQNSHSYSFEILERR |
| Neh2Dual-ACTR$_{10gly78}$-C-DHFR | RDLELPPPYLPSQQDMDLIDILWRQDIDLGVSREVFDFSQRRKEYELEKQKKLEKER<br>QEQLQKEQEKAFFAQLQLDEETGEFLPIQPAQHTQSETSGSGGGGGGGGGGEQVSHG<br>TQNRPLLRNSLDDLVGPPSNLEGQSDERALLDQLHTLLSNTDATGLEEIDRALGIPE<br>LVNQGQALTGCHLEMISLIAALAVDRVIGMENAMPWNLPADLAWFRRNTLNRPVIMG<br>RHTWESIGRPLPGRRNIILSSQPGTDDRVTWVRSVDEAIAAAGDVPEIMVIGGGRVY<br>EQFLPRAQRLYLTHIDAEVEGDTHFPDYEPDDWESVFSEFHDADAQNSHSYSFEILE<br>RR |
